# A neural system that represents the association of odors with rewarded outcomes and promotes behavioral engagement

**DOI:** 10.1101/617902

**Authors:** Marie A. Gadziola, Lucas A. Stetzik, Katherine N. Wright, Adrianna J. Milton, Keiko Arakawa, María del Mar Cortijo, Daniel W. Wesson

## Abstract

Learning strengthens the strong emotional and behavioral responses odors are well known for eliciting. Presumably subserving this, several brain regions display experience-dependent plasticity during odor learning, yet the specific cellular systems involved and the actual influence of these systems on odor-directed behavior are less understood. Here we investigated the transformation of odor information throughout the association of odors with rewards and also sought to link those neural systems with displays of reinforcement-based task engagement. First, we investigated the representation of odor-reward associations within two areas recipient of dense olfactory bulb input, the posterior piriform cortex (pPCX) and olfactory tubercle (OT), using simultaneous multi-site electrophysiological recordings from mice engaged in a reward-based olfactory learning task. As expected, neurons in both regions represented conditioned odors and did so with similar information content, yet both the proportion of neurons recruited by conditioned rewarded odors and the magnitudes and durations of their responses were greater in the OT. Using fiber photometry, we found that OT D1-type dopamine receptor expressing neurons flexibly represent odors based upon reward associations. In both the recordings and imaging, statistically meaningful changes in activity occurred soon after odor onset. Finally, using optogenetics we show that OT D1-receptor expressing neurons strongly influence behavior to promote task engagement. Together our results contribute to a model whereby OT D1 neurons support odor-guided motivated behaviors.

## Introduction

Upon learning, stimuli may acquire meaning which is considered integral in guiding behavioral choice (Berridge and Aldridge, 2008; Chikazoe et al., 2014; Gottfried, 2009; Kumar et al., 2012; Pessoa and Adolphs, 2010; Veldhuizen et al., 2009). Learning to associate a stimulus with an outcome shapes our actions in profound manners – including impacting the choices of foods we eat and can lead to maladaptive outcomes such as those resulting from engaging in compulsive behaviors. This process enables stimuli to hold ‘valence’ (Lewin, 1935) which is of essence in the execution of behaviors that support satiation of basic needs (Lewin, 1935; Tolman, 1932). Despite holding fundamental importance for our day-to-day actions, the neural systems underlying odor associative learning within the brain, that may support the formation of odor valence, are unresolved.

Flexible expressions of neural activity in response to stimuli throughout reward- or emotion-based associative learning are widely observed (*e.g.,* (Berridge, 2019; Gore et al., 2015; Hickey and Peelen, 2015; Isosaka et al., 2015; Morrison and Salzman, 2009; Schoenbaum et al., 1998)). This has been well-established in the orbitofrontal cortex and basolateral amygdala, for instance, wherein neurons in these associative structures display differential firing following reinforcement learning for stimuli conditioned to predict a salient outcome (Roesch et al., 2007; Schoenbaum and Eichenbaum, 1995; Schoenbaum et al., 2003).

Behavioral displays linked to odor associative learning have been observed in all animals studied to date and involve a variety of brain systems (Knaden and Hansson, 2014; Li and Liberles, 2015). Olfactory system structures display forms of plasticity (Barnes et al., 2008; Chapuis and Wilson, 2011; Dias and Ressler, 2014; Doucette et al., 2011; Kass et al., 2013; Lebel et al., 2001; Li et al., 2008; Mandairon and Linster, 2009; Murata et al., 2015; Ross and Fletcher, 2018; Schoenbaum et al., 2000), yet in many of these cases the specific cellular systems involved and/or whether those systems specifically influence behavior are less understood. Among these, secondary olfactory structures including both the piriform cortex (PCX) and olfactory tubercle (OT) may differentially represent odors associated to predict reward availability versus those that do not. Specifically, work from several groups including ours has demonstrated that PCX (Gire et al., 2013a; Roesch et al., 2007; Schoenbaum and Eichenbaum, 1995) and OT neurons (Gadziola et al., 2015; Millman and Murthy, 2020) are more greatly recruited by odors conditioned to predict rewards versus those that are unreinforced. These results have indicated that olfactory structures represent learned odor-reward associations by a combination of recruiting neurons into an ensemble and also divergent firing for rewarded odors.

The representation of odor-reward associations by both PCX and OT neurons leads to the assumption that this function is a distributed, global property among secondary olfactory structures. Indeed, that both PCX and OT neurons encode odor-reward associations with divergent firing (Gadziola et al., 2015; Gire et al., 2013b; Roesch et al., 2007; Schoenbaum and Eichenbaum, 1995) suggests a possible substrate wherein information within these systems is more-or-less equally relayed into downstream structures important for learning and memory (PCX; (Gottfried, 2010; Wilson and Sullivan, 2011)) or those integral for motivated and affective behaviors (OT; (Wesson and Wilson, 2011; Zhang et al., 2017a)). However, an alternative system may be in place: that secondary olfactory structures differentially encode odor-reward associations in manners suggesting that one structure is specialized in value assignment functions. Determining whether these functions are distributed among the PCX and OT is an important question in the overall goal of resolving the neural circuitry underlying odor-guided motivated behaviors.

Based upon the representation of learned odor-reward associations in the OT (Gadziola et al., 2015; Murata et al., 2015), and the positioning of the OT within the ventral striatum (Heimer et al., 1982; De Olmos and Heimer, 1999) wherein it receives dense innervation from dopaminergic neurons in the ventral tegmental area (Ikemoto, 2007; Voorn et al., 1986), we predicted that the OT is a central figure in associating odors with reward contingencies. Infusion of cocaine into the OT is reinforcing, and rodents more robustly seek infusions into the OT than even into the nucleus accumbens (Ikemoto, 2003) – highlighting the likely importance of OT dopamine in influencing motivated states and task engagement, as also supported by a recent optogenetic study (Zhang et al., 2017b). An additional feature of the OT that supports this hypothesis is its cellular composition. The OT is primarily composed of medium spiny neurons expressing either the D1- or D2-type dopamine receptor (Murata et al., 2015). D1-type neurons in many striatal structures are activated by appetitive stimuli and are often considered integral for promoting stimulus reward and approach, whereas at least some reports indicate D2-type neurons are less shaped by reinforcement (for reviews (Lobo and Nestler, 2011; Soares-Cunha et al., 2016). Therefore, it is possible that previous reports of neuromodulation driving reinforcement through the OT (Ikemoto, 2003) is due to direct actions upon OT D1-receptor expressing neurons. Supporting this hypothesis, c-Fos expression is elevated in OT D1-type neurons following associative learning (Murata et al., 2015). Based upon the above, we predicted that OT D1-receptor expressing neurons may support the representation of odor-reward associations in the OT and separately, should therefore also be capable of promoting reinforcer-motivated behaviors, similar to that needed to subserve engagement with odors.

Here we tested the above hypotheses through a combination of multi-site single-unit recordings, cell type-specific fiber photometry, and cell type-specific optogenetic studies, all in mice engaged in reinforcer-motivated operant tasks. All together our results contribute to a model whereby the OT, and OT D1-receptor expressing neurons support odor-guided motivated behaviors.

## Results

### Differential representation of reward-associated odors in the OT versus pPCX

We began by examining whether the representation of odor-reward associations is a global property distributed among the PCX and OT. We monitored OT and pPCX (posterior PCX) single-unit activity from 11 mice implanted with bilateral chronic electrode arrays (**Fig 1A & S1A**) while they engaged in a head-fixed odor discrimination task requiring them to lick a spout for a palatable reinforcer available at the offset of conditioned-rewarded but not unrewarded odors (**Fig 1B-D**). This head-fixed paradigm allows odor-guided behaviors while maintaining precision in odor-delivery (Verhagen et al., 2007). To optimize the data contributed from each animal, all mice were shaped across 6 training phases to discriminate between 2-4 odor pairs (Methods, and **Fig S1B**). Throughout phases 1-4, mice learned a basic lick/no-lick task as we and others have previously described (Gadziola et al., 2015; Verhagen et al., 2007). Phase 5 consisted of pseudo-random trials among which four odors (2 familiar and 2 novel) were separately presented, with the two reinforced odors both resulting in a low-value reward. Finally, in phase 6, the four odors were presented, with one rewarded odor being assigned to a high value saccharin and the other to a low value saccharin. Mice discriminated among the four odors, two of which predicted presentation of reward (either low or high value saccharin), and two unrewarded odors. These sessions include the original training odor set, as well as novel odors presented on a different experimental day (each mouse was shaped on 2 novel sets of four odors on different sessions). Throughout the odor discrimination phases (phases 4-6), performance of the mice was above an 80% correct response threshold (92.6 ± 6.3% (mean ± SD); **Fig S1B**). No major effects were observed between high-versus low-value rewarded odor-evoked responses nor behavior; therefore, herein any odors paired with reinforcers are classified as conditioned rewarded odors, whereas those not paired are termed conditioned unrewarded odors. In previous work from our group (Gadziola and Wesson, 2016), we established that the earliest change in OT firing upon planning to lick is ~500ms prior to lick and thus herein all trials in which animals prematurely licked during the odor were omitted from all analyses of neural activity in order to reduce influences of motor planning or motor execution on data outcomes. Further, we ultimately restrict later analyses to the first 500ms of odor onset to further mitigate influences of lick planning/preparation upon responses.

**Figure 1.**
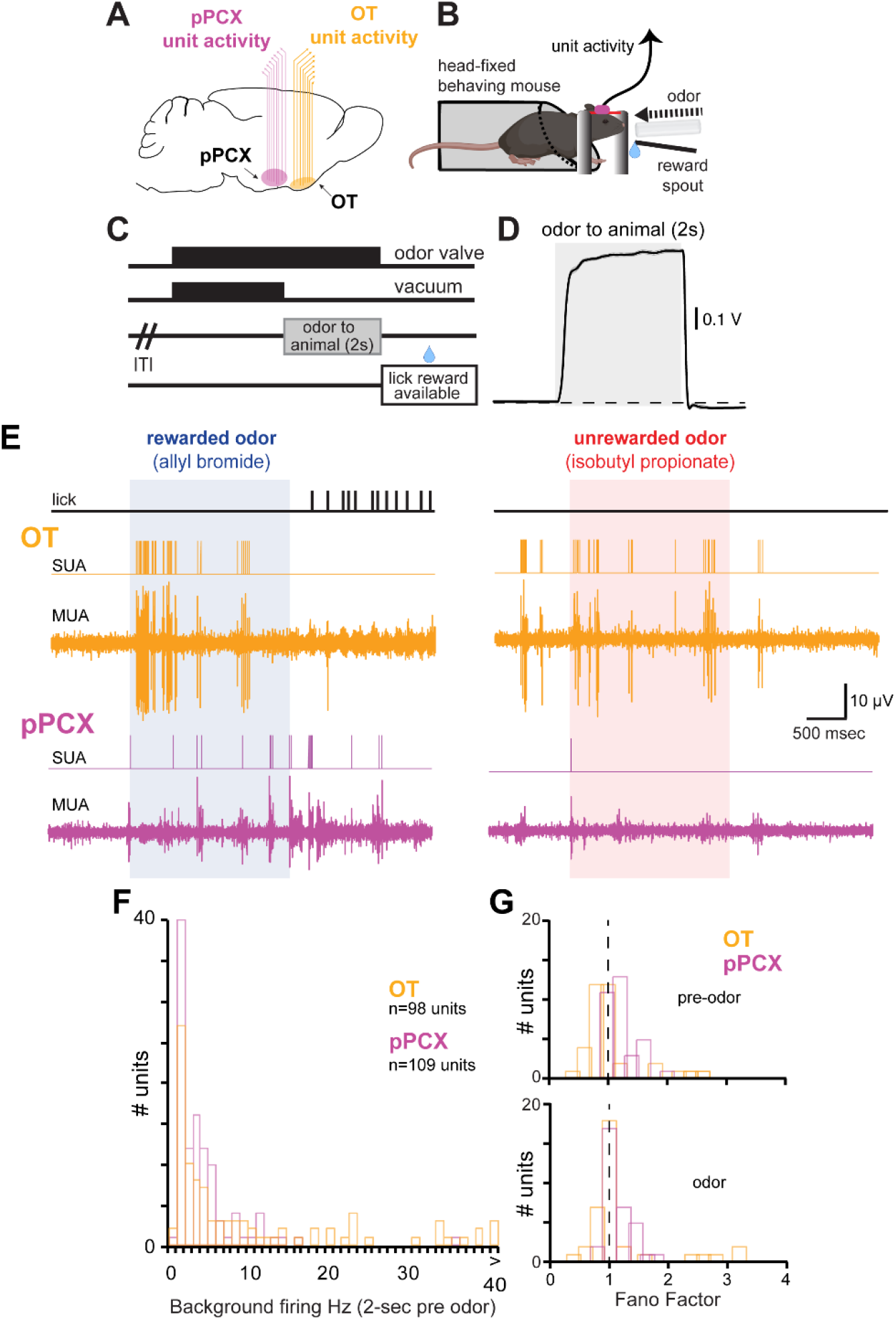
Paradigm for investigating the distribution of odor-reward associations among olfactory cortices. **A)** Recording schematic indicating simultaneous monitoring of OT and pPCX activity with multi-channel arrays in mice (8 wires/region). pPCX and OT activity were recorded on opposite hemispheres within mice. **B)** Schematic of head-fixed behavioral set-up and **C)** experimental trial outline. Water-deprived mice implanted with chronic multi-site arrays into their OT and pPCX (as in A) and a head-bar were acclimated to head restraint and shaped in a lick/no-lick odor discrimination task wherein following a variable inter-trial interval (ITI) one odor signaled the availability of a small palatable fluid reward from a reward spout following odor offset (during the ‘lick reward available’ epoch). Alternative (unreinforced) odors were conditioned to not signal reward availability. OT and pPCX unit activity was acquired throughout performance in the task. **D)** Average evoked PID trace (10Hz low-pass filter) in response to 20 presentations of 1,7-octadiene, illustrating rapid temporal dynamics and good stability of odor delivery within trials. Dashed horizontal line indicates baseline. **E)** Example OT and pPCX unit activity. Multiunit activity (MUA) was spike sorted off-line to identify single-unit activity (SUA) using template matching and cluster cutting with principal component analyses. Vertical lines on the lick trace indicate timing of individual licks (detected with an infrared photo-beam in front of the lick spout). Vertical scale bar applies to both OT and pPCX MUA channels. Data are from two consecutive trials. **F)** Background firing rate distribution of all pPCX and OT single units used for analysis (−2000 to 0 ms pre-odor). *n* = 98 OT units, 109 pPCX units. **G)** Fano factor of OT and pPCX units pre (−2000 to −500 ms; top) and during odor (0 to 1500 ms; bottom). Data from units during discrimination of the original conditioned odors (*viz*., pre-reversal learning; *n* = 36 OT units, 33 pPCX units). Vertical dashed line = the theoretical Fano factor for a Poisson process.

While all 11 mice contributed data from at least one of the brain structures, 7 contributed simultaneous recordings from both the OT and pPCX. Baseline and odor-evoked activity were measured for isolated single neurons. In both structures, odor-evoked firing was observed in response to the odors and occurred soon after odor onset (0.35 ± 0.31s [OT], 0.57 ± 0.48s [pPCX], **Fig 1E**). The baseline firing rates of OT (median rate: 3.1 Hz; range = 0 - 56.9 Hz) and pPCX neurons (median rate: 2.1 Hz; range = 0 - 35.6 Hz) were low across the sampled population (**Fig 1F**). Variability in baseline firing calculated in terms of the autocorrelation function Fano factor (Geisler and Albrecht, 1997; Miller, 2006; Shadlen and Newsome, 1998) was comparable in both structures (Fano factor: OT=1.05 ± 0.2, pPCX=1.18 ± 0.5 (mean ± SD); unpaired *t* test: *t*_(67)_ = 1.45, *p* = 0.15) as was odor-evoked firing in both structures (Fano factor: OT=1.18 ± 0.7, pPCX=1.14 ± 0.2 (mean ± SD); unpaired *t* test: *t*_(67)_ = −0.34, *p* = 0.74) (**Fig 1G**).

As well established, mice learned to discriminate between odor pairs (**Fig S1B**), and following learning we observed that the majority of OT neurons significantly modulated their firing during the task window (65%, 52/80 OT neurons ‘task-responsive’ within 7s of odor onset). An example of one of these OT units across pseudorandom trials of a pair of rewarded and unrewarded odors is displayed in **Figure 2**. Increases in firing upon odor onset is observed across trials of both the rewarded and unrewarded odors (**Fig 2A**), with on average greater firing rates by this OT neuron for the rewarded odor (**Fig 2B**, evident in both raw (top) and Z-score normalized firing histograms (bottom)). The proportion of OT units significantly modulated during the task window was comparable to the population of pPCX neurons modulated in the same time (X^2^(*df*=1, *n*=87) = 3.32, *p*>0.05; 44%, 35/79 neurons). Only these task-responsive neurons were included in subsequent analyses.

**Figure 2.**
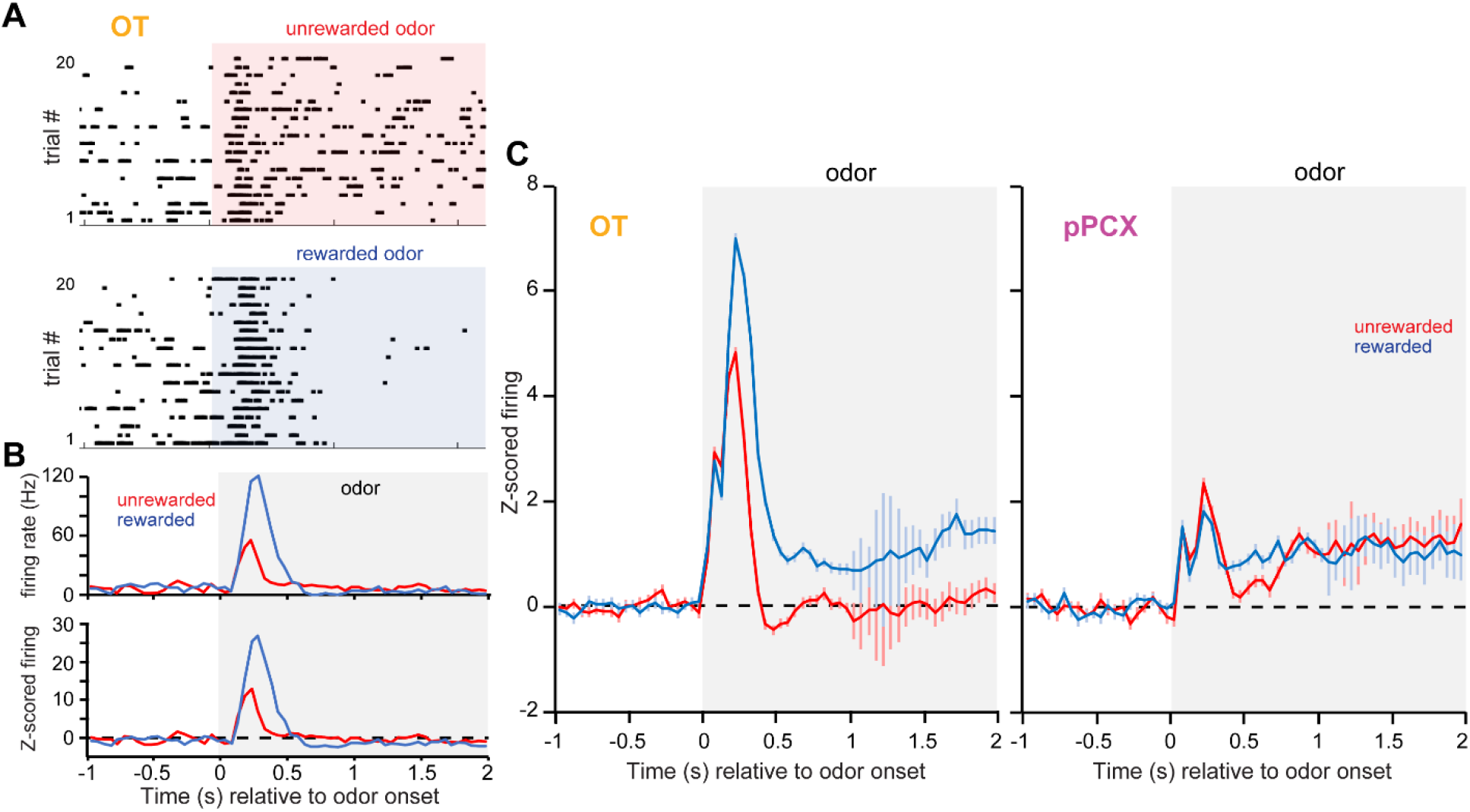
Neural dynamics during odor which reflect reward associations. **A)** Example OT single-neuron rasters in response to a conditioned-rewarded and unrewarded odor. **B)** Average firing rate and Z-score normalized firing (upper and lower panels respectively) calculated across trials for the same neuron in (A). **C)** Peri-stimulus time histograms of Z-score normalized firing rates from task-responsive neurons in the OT (left, n = 104 unit-odor pairs) and pPCX (right, n = 70 unit-odor pairs) from both conditioned rewarded and unrewarded odors. The classifications of ‘rewarded’ and ‘unrewarded’ are global aggregates, with each panel including data from 8-12 different odors. Shaded boxes in all panels indicate the timing of odor.

As expected (Gadziola et al., 2015; Gire et al., 2013b; Millman and Murthy, 2020; Roesch et al., 2007; Schoenbaum and Eichenbaum, 1995), it was common to observe stimulus-evoked changes in firing for both the rewarded and unrewarded odors relative to baseline in both the OT and pPCX (**Fig 2C**). Based upon Z-score, 61.7% (124/201) of OT neuron-odor pairs and 30.9% (42/136) of pPCX neuron-odor pairs displayed significant stimulus-evoked modulation during odor presentation, with significant mean values observed shortly (<500 ms) after odor onset in both regions. Similar to our prior work (Gadziola et al., 2015), OT neurons displayed divergent responses to the odors, with, based on population-averaged Z-score, rewarded odors evoking greater firing rates than those conditioned to be unrewarded (**Fig 2C**). The response to conditioned rewarded odors in the OT included heightened increases in firing within the first 500ms of odor onset, followed by persistence in firing throughout the odor period. In contrast, while pPCX neurons displayed increases in firing to both conditioned odor types, only a brief and modest divergence in responding to both odor types was observed (**Fig 2C**).

Next, we sought to test if this representation of conditioned odors is sufficiently encoded in order to have the potential to be ‘read out’ by downstream targets. We performed decoding analyses (Meyers, 2013) on ensembles of pPCX and OT units sampled from the above population and restricted to only include the original set of odor pairs. Three iterations of the decoding were ran when including randomly sampled populations of either 10, 20, or 30 units from each structure. This analysis revealed that neurons in both regions are able to accurately classify trial response types (hit, miss, false alarm, or correct reject) above chance (**Fig 3A**). The classification accuracy in both regions became greater as the size of the ensemble in both regions increased. No discernable difference between regions was detected in classification accuracy, regardless of the number of neurons included. This was further confirmed when restricting the analyses of classification accuracy to the first 500ms of odor onset, with no statistical differences between regions uncovered in any of the three ensemble sizes (two-sample *t* tests, *p* > 0.05; **Fig 3B**). Thus, both OT and pPCX ensembles are capable of accurately classifying trial type during reward-motivated odor discriminations, indicating that at the ensemble level, each region provides similar information content.

**Figure 3.**
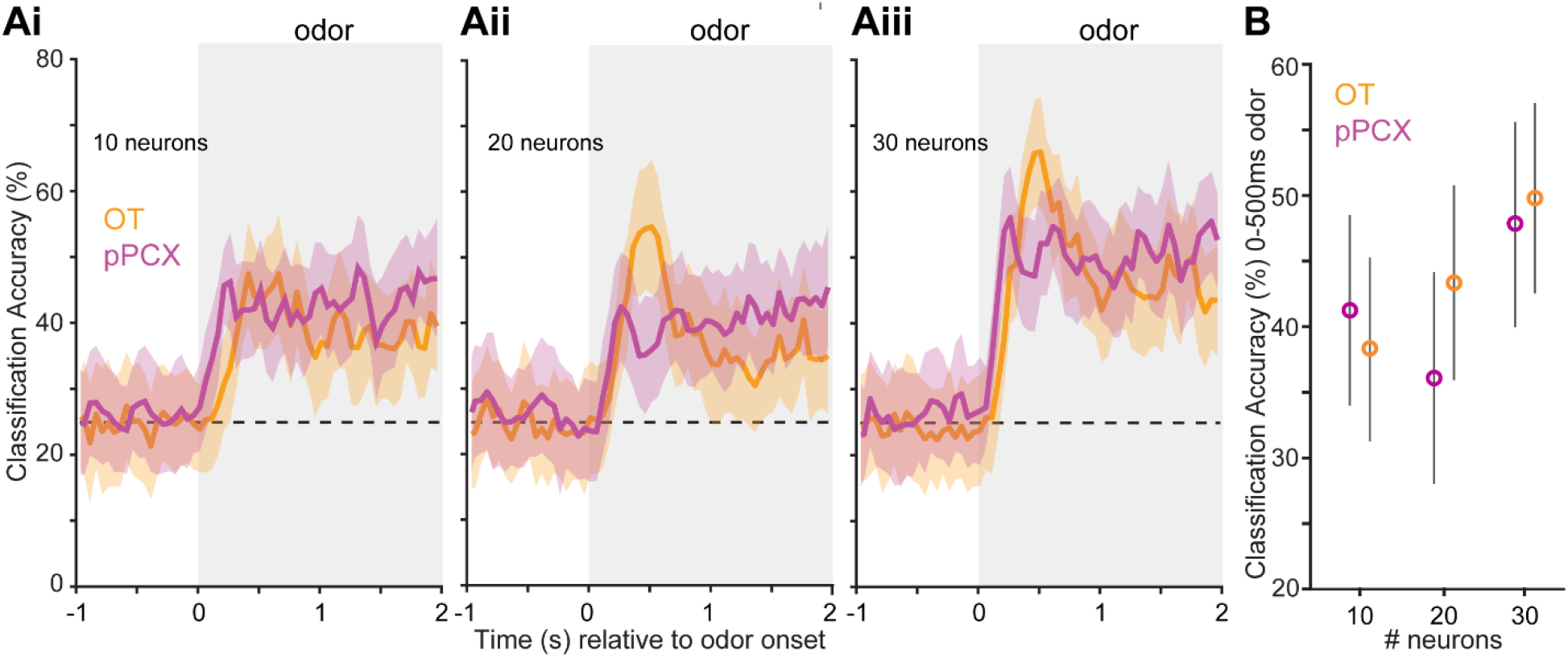
Classification accuracy for pPCX and OT units to differentiate trial type. **A)** Peri-stimulus time histograms of accuracy for classifying trial types (conditioned rewarded versus unrewarded). Ensembles from both brain regions display classification accuracy greater than chance (probability line = horizontal dashed line) and above the shuffled data beginning soon after odor onset which while not displayed for visual simplicity never exceeded 33.7. Accuracy became greater with increasing numbers of randomly sampled neurons provided to the classifier (10 neurons to 30 neurons, **Ai** to **Aiii** respectively) which was also observed when the analysis window was restricted to the first 500ms of odor onset **(B)**. Shaded boxes indicate the timing of odor. Values represent the mean ± SD of the resamples.

As the population-based analyses captured all units regardless of response type (odor-excited vs suppressed), we next examined the influence that response type had in each region, looking both at the population-level responses and individual neurons, along with their time courses of significant responding (**Fig 4**). Of the neuron-odor pairs that displayed significant stimulus-evoked modulation during odor presentation (those defined in **Fig 2C**), the vast majority of those neurons showed increases in activity relative to baseline, with the remaining neurons being suppressed. The proportion of odor-excited neurons were highly similar between regions (84.7% of OT and 85.7% of pPCX). The temporal response pattern of neurons that increased vs decreased activity relative to baseline in both structures indicated they each did so most greatly within the first 500 ms of the odors (**Fig 4A**). Knowing this, to test whether individual OT and pPCX neurons preferentially respond to one conditioned odor over the other, we analyzed the absolute Z-score values within individual neurons, computed as an average across the first 500ms following odor onset. Individual OT neurons preferentially responded to rewarded odor with greater absolute Z-score values than the unrewarded odors (4.21 ± 0.4 vs 2.81 ± 0.29, respectively; paired *t* test: *t*_(199)_ = −2.95, *p* = 0.003), whereas pPCX neurons did not respond differentially to rewarded and unrewarded odors across the population (1.85 ± 0.23 vs 2.06 ± 0.26, respectively; paired *t* test: *t*_(134)_ = 0.45, *p* = 0.65). Furthermore, averaged absolute Z-score values in response to rewarded odors were greater for the population of OT neurons when compared to pPCX neurons (4.21 ± 0.4 vs 1.85 ± 0.23, respectively; two-sample *t* test: *t*_(165)_ = −4.45, *p* < 0.001).

**Figure 4.**
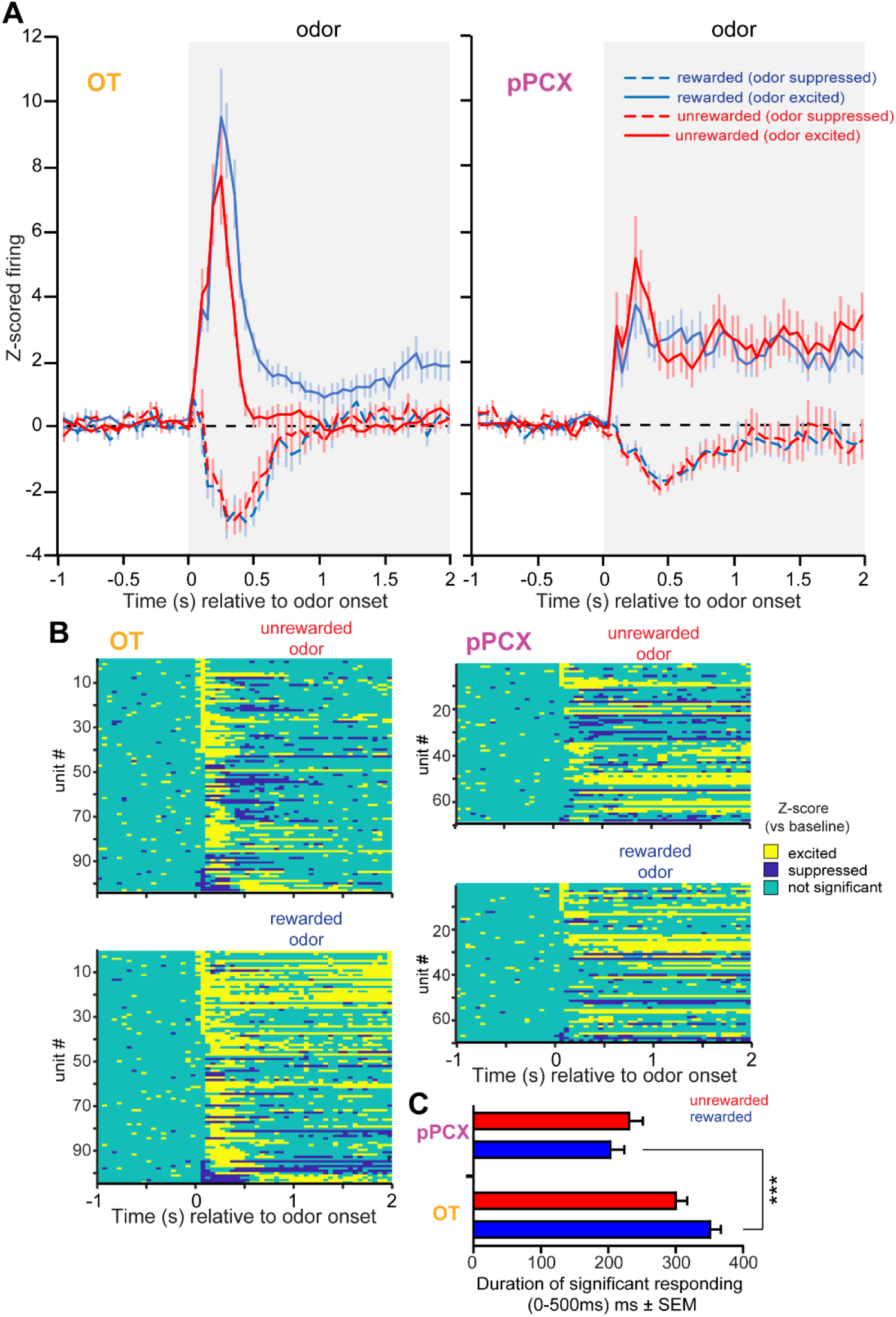
Differential representation of reward-associated odors in the OT compared to the pPCX. **A)** Peri-stimulus time histograms of Z-score normalized firing rates in the OT (left) and pPCX (right) from both conditioned rewarded and unrewarded odors as in **Figure 2** but separated by whether the response was excited or suppressed relative to baseline. Population values represent the mean ± SEM. Shaded boxes indicate the timing of odor. **B)** Z-score results from individual neuron-odor pairs indicating timing of significant bins. Data are organized in descending order based upon the significant excitation in the first 50ms bin. **C)** Cumulative time significantly responding neurons were modulated during the first 500ms of odor onset, on average across neurons (mean ± SEM). ***=*p*<0.001. *n* = 100 OT, 68 pPCX neuron-odor pairs. See Results for additional statistical outcomes.

We additionally compared the size of the population modulated by reward-associated odors and its duration of significant Z-score responses in the first 500 ms after odor onset across the population of neurons in both regions (including units with ≥ 1 significant bin) (**Fig 4B**). More neurons were modulated by reward-associated odors in the OT (99%, 99/100) than the pPCX (85%, 58/68) (X^2^(*df*=1) = 10.287, *p*=0.0013; *n* = 168 total neuron-odor pairs). Further, OT responses were significantly longer in duration compared to pPCX responses (328 ± 11 vs 219 ± 13 ms, respectively; two-sample *t* test: *t*_(335)_ = −6.51, *p* < 0.001). OT response durations to rewarded odors were longer than unrewarded odors (354 ± 15 vs 303 ± 14 ms, respectively; paired *t* test: *t*_(99)_ = 2.70, *p* = 0.008), and longer than pPCX responses to rewarded odors (**Fig 4C**) (354 ± 15 vs 206 ± 19 ms, respectively; two-sample *t* test: *t*_(166)_ = −6.26, *p* < 0.0001). Together, these analyses including 1) decoding accuracy, 2) the proportion of modulated units, 3) their Z-score magnitudes, and 4) their durations of significant responses, indicate that neurons in both regions represent conditioned odors and do so with similar information content, yet that both the proportion of neurons recruited by conditioned rewarded odors and the magnitudes and durations of their responses were greater in the OT.

### D1 receptor-expressing OT neurons display divergent responses to conditioned rewarded odors

Which neurons are responsible for the pronounced display of reward-associated odor representations in the OT? We addressed this question by performing fiber photometry-based imaging (*e.g.,* (Gunaydin et al., 2014; Kudo et al., 1992)) of GCaMP6f (Chen et al., 2013) from D1 receptor-expressing neurons in the OT which we predicted may differentially represent reward-associated odors. We also imaged D2 receptor-expressing neurons in the OT of separate mice as comparison. Mice expressing Cre-recombinase in either neurons expressing the D1 receptor (D1-Cre) or D2 receptor (D2-Cre) (Gong et al., 2003) were injected with floxed GCaMP6f into their OT and later implanted with optical fibers (see Methods, **Fig S2A**) before being water deprived and shaped in phases 1-4 of the lick/no-lick odor discrimination task used above. During imaging (**Fig 5B**), the GCaMP6f emission spectrum was collected as a measure of aggregate OT D1 or D2 neuron activity. Simultaneously, from all recordings, we also collected endogenous UV emission spectrum as a control measure for movement artifacts (**Fig S2B**). The resultant GCaMP6f signal was subtracted from the UV signal for a single output, which is considered closely reflective of aggregate neural activity.

**Figure 5.**
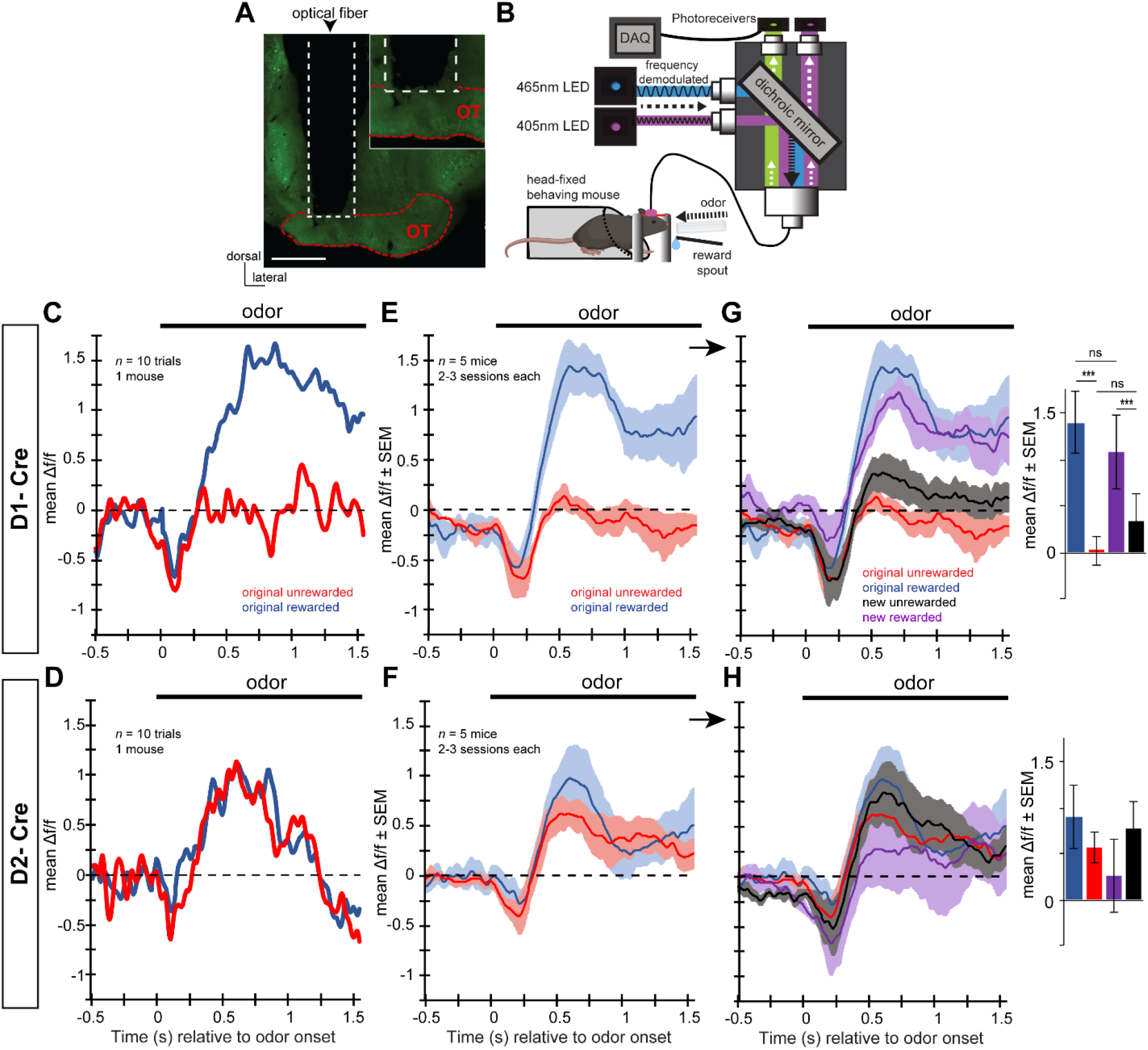
OT D1 receptor expressing neurons are differentially modulated by reward-associated odors. **A)** Example localization of an optical fiber implant into a mouse injected with GCamP6f (green). The optical fiber was positioned to terminate immediately dorsal to the OT. **B)** Schematic of the fiber photometry system for use in head-fixed behaving mice. 465nm and 405nm LEDs excitation wavelengths passed through dichroic mirrors prior to being sent into the implanted optical fiber via a patch cable. GCaMP6f and UF emission were then both amplified by femtowatt photoreceivers prior to being digitized simultaneously with behavioral data and stimulus presentation events. Please see Fig 1 for a description of the head-fixed paradigm. Examples of separate GCaMP6f and UV emission can be found in **Supplemental Fig 2B**. Example GCaMP6f responses to the original rewarded and unrewarded odors in a single D1-Cre **(C)** and D2-Cre mouse **(D)** averaged over 10 trials of correct behavioral responses during blocks of criterion behavioral performance. These example averaged traces indicate divergent responses to the original rewarded odor vs the original unrewarded odor in D1-Cre but not D2-Cre mice, which is maintained at the population level when averaging across 5 mice, throughout 2-3 sessions during criterion behavioral performance (**E & F**). The encoding of odor-reward associations in D1-Cre mice (as in **E**) was confirmed in experiments wherein the odor-reward contingencies were reversed (original rewarded = no unrewarded, and *vice versa*) (**G & H**). Insets in **G & H** display the mean delta f/f during 0.5-0.75sec of odor in for all four stimulus types. ***=*p*<0.001. ns = not significant (*p*>0.05). No significance was found in any planned comparison in the D2-Cre mice (**H**).

We imaged 5 D1- and 5 D2-Cre mice with confirmed fiber implants and GCaMP6f expression both in the OT (**Fig S2A**) throughout learning and performance in the lick/no-lick odor discrimination task (**Fig S2C & S2D**). To enhance each animal’s contributions for a more rigorous data set, we shaped each mouse on several odor pairs over differing daily sessions, yet only one pair of odors in a given session. This allowed each animal to contribute odor-modulated GCaMP6f signals to 2-3 rewarded and unrewarded odors (on different sessions) at or above criterion performance (mean rewarded odor trials / session = 44.5 [24-75 min-max range]; mean unrewarded odor trials / session = 91.3 [52-133 min-max range]). In these experiments mice tended to lick prematurely and thus while there are a sufficient number of both odor trial types, there are less rewarded odor trials. Across all rewarded and unrewarded odors, we observed modulations in both D1 and D2 neuron activity following odor onset. This was evident both looking within individual mice at subsets of trials (**Fig 5C & D**), and averaged across mice over several sessions (**Fig 5E & F**). No discernable differences were observed in the time prior to odor onset, or before 0.5s following odor onset between any stimulus type in either D1 or D2 mice (**Fig 5E & F**). Later during the odor period however (0.5-0.75s), across all D1 mice, conditioned rewarded and unrewarded odors elicited differing amounts of activity ((*F*(3,19)=15.002, *p*<0.001); **Fig 5E**). Whereas D1 neurons were largely unmodulated by the conditioned unrewarded odor, conditioned rewarded odors elicited large increases in activity during the odor. In contrast, no differential representation of conditioned rewarded versus unrewarded odors was detected in D2 mice, with both conditioned stimuli eliciting similar increases in activity during odor (*F*(3,19)=0.84, *p*=0.493); **Fig 5F**).

### The responses of D1 receptor-expressing OT neurons are flexible and encode the associated reward outcomes of odors

Having imaged OT neurons and observed that D1-type dopamine receptor expressing neurons represent odor-reward associations, we were next able to ask whether this differential representation of conditioned rewarded odors observed among the D1 neuron population is flexible. In other words, does the representation of conditioned-rewarded odors flexibly follow as odors are associated to predict new outcomes? To address this, in a separate behavioral session we used reversal learning to reverse the odor-outcome contingencies so that a previously rewarded odor is no longer rewarded and a previously unrewarded odor is now rewarded. Most mice were shaped upon at least 2 odor-reversals (differing odor pairs, range 2-3). If a population of neurons is ‘identity’ encoding they should exhibit the same odor selectivity before and after reversal learning, whereas a population encoding the reward outcome of an odor (rewarded or not) should reverse its odor preference (Gire et al., 2013b; Roesch et al., 2007). Animals took several sessions to learn the reversal to criterion in this task requiring them to withhold licking until odor offset (**Fig S2C and S2D**), and comparisons were made only from sessions following task acquisition, during blocks consisting of criterion-level performance, and correct responses during each trial.

On average across animals, the evoked GCaMP6f signals in D1-Cre mice tracked the behavioral contingencies. Pairwise comparisons of the means using least significance difference testing revealed that the original rewarded odor and the newly rewarded odor eliciting evoked responses with similar mean amplitudes (*1(9), p=*0.202; **Fig 5G**). Likewise, the original unrewarded odors elicited similar responses as the new unrewarded odors (*1(9), p=*0.177). In contrast, no effect of reversal on either the rewarded (*1(9), p=*0.785) or unrewarded odors (*1(9), p=*0.522) were observed in the D2-Cre mice (**Fig 5H**). D2 neurons maintained somewhat positive responses for all odors, regardless of reward association. These data support our hypothesis that D1 receptor-expressing neurons are responsible for the profound representation of odor-reward associations in the OT.

### OT D1 receptor-expressing neurons support behavioral engagement

Having observed reward-associated odor coding in OT D1 receptor-expressing neurons, we next asked whether the activity of these neurons influences engagement and execution of motivated behaviors. Indeed, the latency and magnitude whereby OT D1 neurons divergently represent conditioned rewarded vs unrewarded odors may serve as an effective way whereby to influence motivated behavior. To test this, D1-Cre mice injected with a floxed AAV encoding channelrhodopsin (ChR2) [n = 7] (or floxed AAV encoding a reporter fluorophore [n = 5]) into their ventral striatum were later implanted with an unilateral optical fiber into their OT (**Figs 6A & B**). While ChR2 expression may not be selective to the OT following injection, the fiber-optic tips for stimulation of ChR2 were localized in the OT (**Fig 6A**).

**Figure 6.**
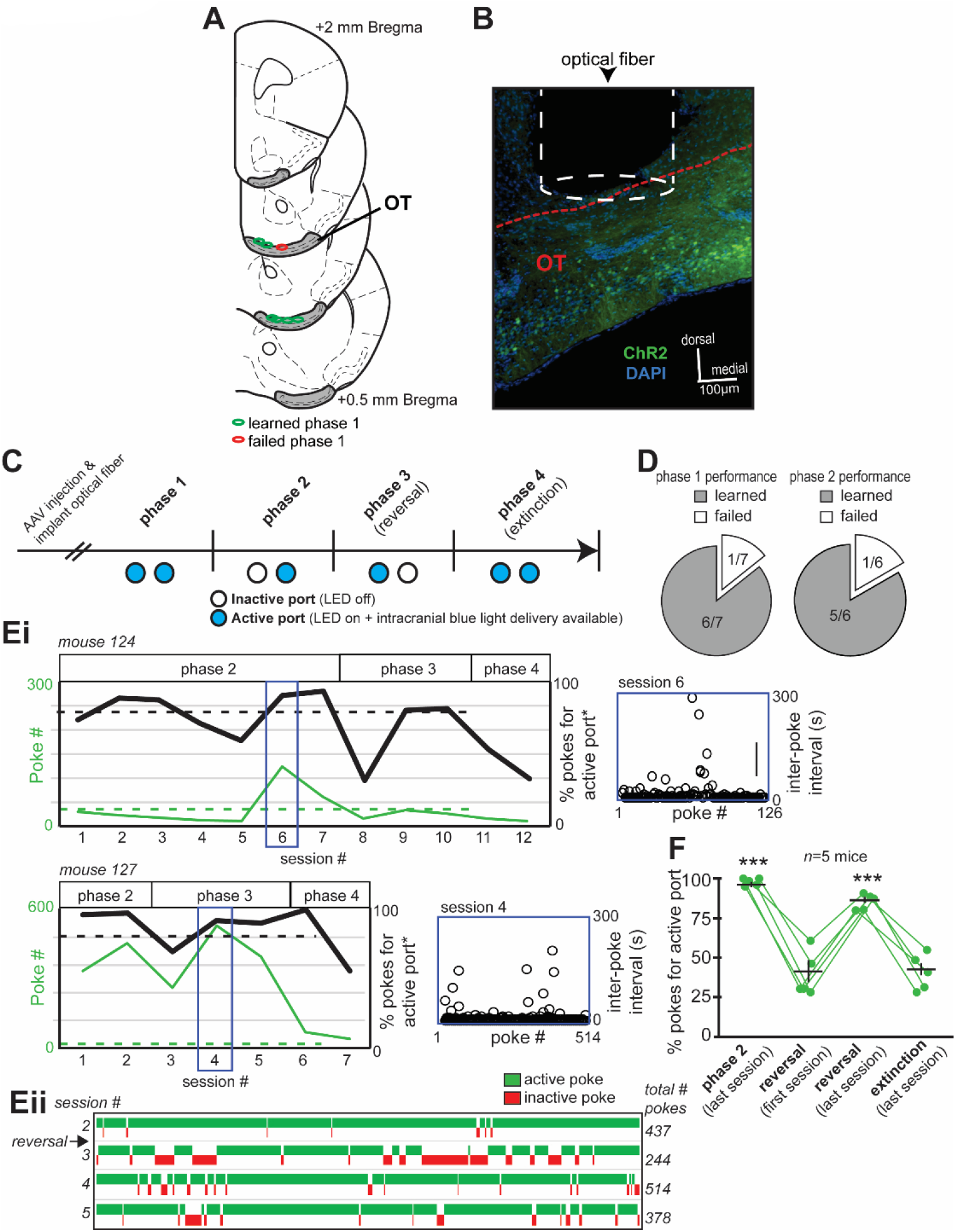
OT neurons expressing the D1 receptor promote behavioral engagement. **(A)** Optical fiber implant locations from mice contributing optical-intracranial self-stimulation data (opto-ICSS), segregated based upon those mice that learned versus failed to acquire phase 1 of the task. **B)** Representative image of an optical fiber positioned within/immediately above the OT of a D1-Cre mouse that was previously injected with AAV.ChR2. Red dashed line indicates the dorsal border of the OT. **(C)** Opto-ICSS task schematic wherein mice were required to nose poke in exchange for blue-light mediated optogenetic excitation of OT D1 neurons. At least 2 weeks following injection with either a floxed AAV encoding ChR2 and a reporter fluorophore (AAV.ChR2) or a floxed AAV solely encoding a reporter fluorophore (AAV), mice were shaped on the opto-ICSS task. Please see text for a description of the four task phases. **(D)** Pie chart indicating that all but one AAV.ChR2 mouse reached criterion performance on phase 1 of the task. **(Ei)** Performance of two example mice during phases 2-4 of the opto-ICSS task. The mice displayed diversity in their number of pokes into the active port, and in their % of pokes displayed for the active (blue-light emitting) versus inactive ports. Both of these mice reached criterion on phase 2 and phase 3 wherein they had to learn to redirect their poking for a new active port location (reversal learning). Shown also is the behavior of these two mice during extinction, wherein the port lights were on (*viz.,* both ports were visually ‘active’) but no optogenetic stimulation was available regardless of poking. *data plotted as ‘active port’ referring to the previously active port in phase 3. Blue insets illustrate each animal’s inter-poke intervals (open circles) throughout the indicated behavioral session. **(Eii)** A win-loss plot for mouse 127 from session 2-5. **(F)** Quantification of opto-ICSS data indicating that ChR2 mediated stimulation of OT D1 neurons promotes task engagement. The first phase of reversal learning resulted in a significant reduction in % pokes for the new active port, which with experience was restored on the last session of the reversal phase, and again reduced during subsequent extinction testing. Data points indicate individual mice. \*\*\**p*<0.0001.

Implanted mice were shaped in an optical intra-cranial self-stimulation task, adapted from (Carlezon and Chartoff, 2007; Ilango et al., 2014; Vicente et al., 2016), allowing for assaying of stimulation-seeking goal-directed task engagement (**Fig 6C**). Shaping in this instrumental task occurred over several phases (See Methods). In the first phase, mice could poke their snout into either of two ports in exchange for blue light stimulation into their OT (25Hz, for 2s, fixed ratio 1). All but one of the D1-Cre mice injected with AAV.ChR2 reached a set level of criterion performance (40 pokes into either port during a 1 hr session) (**Fig 6D**), indicating that they readily learned to poke in exchange for blue light stimulation. The single AAV.ChR2 injected mouse that did not reach criterion only poked 12 times in the last session. No control AAV injected mice reached criterion performance, with the maximum poke numbers for each mouse within sessions ranging from 1-27 pokes. The mice reaching criterion on Phase 1 then progressed onto Phase 2 wherein only one of the two ports delivered light. The port receiving the greater number of pokes during Phase 1 was selected as the ‘active’ port for Phase 2. The vast majority of AAV.ChR2 mice (5/6, **Fig 6D**) reached criterion on Phase 2 and example performance of two of these mice is shown in **Figure 6E**. To ensure that these mice were indeed seeking light stimulation, on the subsequent Phase 3, the location of the active port was reversed and despite this, all 5 mice who reached criterion on Phase 2 redirected their responding to criterion levels for the new active port. Compared to the first session of reversal phase testing, there were significantly more pokes for the active port during both the last session of phase 2 testing (paired *t* test: *t*_(4)_ = 7.62, *p* = 0.0016), as well as during the last session of reversal (paired *t* test: *t*_(4)_ = −6.84, *p* = 0.0024) (**Fig 6F**). This finding that the ChR2 injected D1-Cre mice ‘followed’ the active port upon reversal indicates that operant responding in these mice is neither simply a product of implant laterality (which we varied across mice, see Methods) or simply due to lateralized port preferences. Finally, in Phase 4 (‘extinction’) we attempted to extinguish responding by rending both ports inactive over two sessions which resulted in the percentage of pokes for the previously active port as well as the number of pokes overall in both ports to be reduced compared to the last session of Phase 3 testing (**Fig 6Ei & 6F**). Mice displayed significantly fewer pokes for the active port on the last extinction session than compared to the last session of phase 2 testing (paired *t* test: *t*_(4)_ = 14.88, *p* < 0.0001), as well as during the last session of reversal (paired *t* test: *t*_(4)_ = 14.38, *p* < 0.0001) (**Fig 6F**).

The maximum total number of pokes displayed by each mouse within a given session of phase 2 or 3 performance varied considerably, with one mouse poking 514 times yet another only 56 times (across animal range 226.8 ± 191.5 [mean ± SD]). We used the same sessions contributing the maximum poke numbers as above to calculate the duration of time in between pokes as an index of engagement throughout the session. Not surprising based upon the variance in poke numbers across mice, so too did inter-poke intervals vary across mice, with one mouse displaying a mean interval as rapid as 6.7 ± 17.2 sec (across animal range 30.45 ± 33.8 sec [mean ± SD]). Notably, within animals, the inter-poke intervals were mostly stable within the given session which suggests mostly sustained task engagement (**Fig 6Ei**, blue insets). These results indicate that OT neurons expressing the D1-type dopamine receptor support task engagement.

## Discussion

Our sensory systems hold a remarkable capacity for experience-based plasticity and informing a wide-range of critical behaviors. Olfactory structures are well-known for their plasticity (Barnes et al., 2008; Chapuis and Wilson, 2011; Dias and Ressler, 2014; Doucette et al., 2011; Kass et al., 2013; Lebel et al., 2001; Li et al., 2008; Mandairon and Linster, 2009; Murata et al., 2015; Ross and Fletcher, 2018; Schoenbaum et al., 2000), yet in many of these cases the cellular systems involved and/or whether those systems specifically influence behavior are less understood. Attraction to odors may employ innate (unlearned) or acquired (learned) mechanisms. Since the olfactory bulb is the starting-point for odor perception, it is not surprising that the olfactory bulb holds importance for both innate and learned odor hedonics (*e.g.,* (Doucette et al., 2011; Kermen et al., 2016; Kobayakawa et al., 2007; Wilson et al., 1987). Indeed, recent work uncovered evidence that a specific zone within the main olfactory bulb is necessary for generating odor hedonics (Kermen et al., 2016). The hedonic information of odors in the OB may then be distributed into the OT and PCX. Both PCX and OT neurons may be recruited to encode odors associated with rewards and do so with divergent firing to conditioned rewarded versus unrewarded odors (Gadziola et al., 2015; Gire et al., 2013b; Roesch et al., 2007; Schoenbaum and Eichenbaum, 1995). The major goal of this work was to understand what each of these brain regions may contribute to behaviors driven through the association of odors with rewards, and what cellular systems are involved in these behaviors.

### The way OT units represent odor-reward associations is unlike that in the pPCX

A key advance the present study yields is that while both OT and pPCX ensembles contain similar amounts of information regarding the learned odors (*e.g.,* **Fig 3**), they differ significantly in the manner by which they represent these odors. This includes differences in the proportion of modulated units, their Z-score magnitudes, and their durations of significant responses. Specifically, the proportion of OT units significantly representing reward associated odors far surpassed the proportion of pPCX units, with greater response magnitude and longer duration of modulation for conditioned rewarded odors observed in these OT units relative to pPCX units (**Fig 4**). Our findings have both similarities and differences to the recent work of Millman and Murthy who employed comparable OT and pPCX unit recordings to also uncover differential representations of odor-reward associations in mice (Millman and Murthy, 2020). In terms of differences, Millman and Murthy reported that pPCX neurons represent odor identity but do not represent odor-reward associations, whereas, consistent with work by other groups (Calu et al., 2007; Gire et al., 2013b; Roesch et al., 2007; Schoenbaum and Eichenbaum, 1995), our analyses did (**Fig 3**). Another difference is that Millman and Murthy reported OT units to be highly variable in their responding, yet our analyses indicated considerable stability (**Fig 1G**). Both of these differences could be due to differences in behavioral shaping or subtle nuances in analyses between the present study and theirs. In terms of similarities, Millman and Murthy’s work also showed that OT neurons profoundly represent odor-reward associations. Our group was the first to report that odor identity and odor-reward associations (is it rewarded or not?) may be multiplexed in some OT neurons (Gadziola et al., 2015). We built off of that to show that OT neurons even represent instrumental responding, reward sizes, and types, and do so depending upon motivational levels (Gadziola and Wesson, 2016). Just like in our original studies, Millman and Murthy reported that odor-reward associations influence OT neuron activity within just a few hundred ms after odor. Interestingly, the Millman and Murthy paper highlights the tendency for OT neurons to rapidly form odor-reward associations throughout learning which had not been carefully examined before. Taken together, all of these results highlight the likely importance of the OT in odor-motivated behaviors (Gadziola et al., 2015; Millman and Murthy, 2020; Murata et al., 2015; Zhang et al., 2017b).

The inter-regional differences we observed in how neurons represent odor-reward associations could not simply be the result of behavioral, affective, or other state-dependent influences since recordings were largely sampled from both the pPCX and OT simultaneously. In our task, animals would not be reinforced if they licked during the odor period and any trials wherein mice did so were discarded from analysis. While we cannot rule out that planning to lick does not influence the outcomes of the present study, in our previous work (Gadziola and Wesson, 2016), we found that OT neurons would only display firing for several hundred ms prior to a lick. By restricting the analyses of both the unit and photometry data herein to less than the first second of odor in both data sets, we sought to minimize possible influences of motor planning/lick preparation in outcomes, although these factors still may be of consequence. Notably, some aspects of the temporal dynamics whereby OT neurons were modulated by rewarded odors differs from that observed in the pPCX. The response to conditioned rewarded odors in the OT included heightened increases in firing within the first 500 ms of odor, followed by persistence in firing throughout the rest of the odor period that diverged from firing in response to unrewarded odors. In contrast, while pPCX neurons displayed increases in firing to both conditioned odor types, only a brief and modest divergence in responding to both of the odor types was observed (**Fig 2C**). Whether this interesting dynamic may aid in signaling to an animal that the odor is or is not predictive of reward availability (*viz.*, ‘lick’ or ‘do not lick’) will be an interesting question for future studies. Indeed, whether and how the observed neural dynamics may in fact inform behavioral choice will need future investigation. Overall, this work uncovers that systems downstream from the OB are specialized in their representation of odors which predict reward availability.

### OT D1 neurons flexibly represent odor-reward associations

We found that OT D1-type dopamine receptor expressing neurons represent odor-reward associations. This significantly extends previous work in awake animals recording OT units of unknown identity (Carlson et al., 2018; Gadziola and Wesson, 2016; Gadziola et al., 2015; Millman and Murthy, 2020) by indicating what OT cell type may be responsible for encoding odor-reward associations. We found OT D1-type dopamine receptor expressing neurons not only differently represented rewarded versus unrewarded odors, but that they did so flexibly throughout reversal learning. An exciting preprint showed that midbrain phasic dopamine release into the OT may be sufficient for odor-reward associations among OT single units (Oettl et al., 2019). That outcome, plus work showing the influence of OT dopamine on odor preferences (DiBenedictis et al., 2014; Zhang et al., 2017b), makes it highly likely that dopamine acts upon OT D1 neurons in manners integral to behavioral displays of odor valence.

There are some notable differences between the observed dynamics of the OT single-units and OT D1 neural activity worthy of discussion. For instance, in the fiber photometry of D1 neurons we observed transient suppression below baseline followed by excitation for conditioned rewarded odors which surpassed that of conditioned unrewarded odors. The transient suppression was not observed among the OT single-units which suggests some difference in either the subtle behavior of the animals in these two separate preparations (performed on different set-ups and by different experimenters), or in the origins of the signals themselves. Both are certainly possible to consider as caveats in reconciling the dynamics yielded by two preparations. While our single-unit recordings likely provided ‘unbiased’ monitoring of various neurons throughout the OT, not targeted to any specific neuron type, it is possible some dynamics are not detected. In the fiber photometry preparation, not only are we monitoring ‘aggregate’ neural activity, but also some GCaMP was detected outside of the OT, including in some PCX neurons (*e.g.,* **Fig 5A**). While PCX fibers innervate the OT and influence its activity (White et al., 2019), it is unlikely that these fibers contributed greatly to our measured photometry responses in contrast to the OT neurons themselves. Future work to understand how population-level GCaMP responses reflect that of populations of diverse individual neurons is needed to resolve this.

### The influence of OT D1 neurons on engagement

Our results support a role for OT D1 neurons, not unlike D1 neurons in the nucleus accumbens (Lobo and Nestler, 2011), in behaviors integral for valence -- reinforcement learning and task-engagement. To come to this conclusion, we used a straightforward instrumental responding paradigm wherein mice were allowed to nose-poke in exchange for OT D1 neuron stimulation. This paradigm has several advantages over real-time place preference testing or other tests that serve to assay motivated behaviors since it allows for measures of discrete, stimulus (*i.e.*, light) seeking behaviors (nose-pokes). The finding that some animals responded at the rate they did, indeed up to 500 pokes in an hour, provides a compelling demonstration that OT D1 neurons influence motivation states, or tap into reward circuitry, in a manner which promotes task engagement. This test was done in the absence of any experimentally-delivered odors which further reinforces the notion that optical stimulation of OT D1 neurons engages important internal circuitry for guiding behavior, versus that which may get paired with odor information entering the OT from the olfactory bulb. OT D1-receptor expressing neurons send efferents throughout the brain’s affective and sensory centers (Zhang et al., 2017a). OT D1 neurons may signal to downstream systems that also influence motivation like the amygdala or hypothalamic nuclei. This model, as also supported by immediate early gene mapping experiments following olfactory learning (Murata et al., 2015), would be potentially powerful in cases wherein odors associated with positive outcomes, which as we show here are represented in OT D1 neurons, could influence downstream limbic systems to strengthen behavioral responding through their coordinated output. What specific targets of OT D1 cells are responsible for behavioral outcomes will be an important question for future research to address. It is notable that striatal neurons, including medium spiny neurons, extend collaterals onto one another which will result in modulation of neighboring cells (Taverna et al., 2008), and thus D2 neurons may have a role in influencing D1 neuron spontaneous and stimulus-evoked activities.

### Conclusions

Taken together, our results contribute to a model whereby the OT is a specialized center for encoding odor-reward associations in manners which may be important for informing stimulus valence and also possibly upcoming actions. This along with the finding that OT D1 neurons encode reward-associated odors and motivate reinforcement-seeking in a nose-poke based task which emulates snout-directed investigation behaviors like those used by rodents to sample odors, leads us to predict that OT D1-receptor expressing neurons are a fundamental component of brain systems needed to inform and respond to odors.

## Supporting information

Supplemental Results

## Acknowledgements

This work was supported by NIH grants R01DC014443, R01DC016519, R01DA049545, and R01DA049449 to D.W. Some mouse strains used in this research were obtained from the Mutant Mouse Regional Resource Center (MMRRC), an NIH funded strain repository, and were donated to the MMRRC by the MMRRC facility at the University of California, Davis.

## Author Contributions

Conceptualization: M.A.G., L.A.S., D.W.W.; Methodology: M.A.G., L.A.S., D.W.W.; Investigation: M.A.G., L.A.S., K.N.W., A.J.M., K.A., M.C.S., D.W.W.; Writing – Original Draft: M.A.G., L.A.S., D.W.W.; Writing – Review & Editing: M.A.G., K.N.W., D.W.W.; Visualization: M.A.G., L.A.S., D.W.W.; Supervision: D.W.W.; Funding Acquisition: D.W.W.

## Methods

### LEAD CONTACT AND MATERIALS AVAILABILITY

Further information and/or requests for resources, code, or other materials should be directed to the Lead Contact, Dan Wesson, at. This study did not generate any new animal models nor reagents.

### EXPERIMENTAL MODEL AND SUBJECT DETAILS

Eleven 2-4 months of age male C57bl/6 mice originating from Harlan Laboratories (Indianapolis, IN) were used for single-unit recordings. For fiber photometry and optical intra-cranial self-stimulation, we utilized similar aged (2-4 months) male bacterial artificial chromosome transgenic mice expressing Cre-recombinase under control of the *drd1a* (D1-Cre; obtained from the UC Davis Mutant Mouse Regional Resource Center, strain EY262Gsat/Mmucd) or *drd2* gene (D2-Cre; obtained from the UC Davis Mutant Mouse Regional Resource Center, strain ER44Gsat)(Gong et al., 2003). Genotyping was performed following standard protocols using tail tissue DNA. Mice were housed on a 12:12 h light-dark cycle with food and water available *ad libitum*, except when water was restricted for behavioral training (see below). Up to 5 mice were co-housed in a cage before experimentation, but all post-surgical animals with any cranial implants were housed individually. All experimental procedures were conducted in accordance with the guidelines of the National Institutes of Health and were approved by the Case Western Reserve University and University of Florida Institutional Animal Care and Use Committees.

## METHOD DETAILS

### Methods for multi-site single-unit recordings during lick/no-lick odor discriminations

#### Intra-cranial electrode implant surgery

The electrode implants were conducted as described previously (Gadziola et al., 2015; Gadziola & Wesson, 2016). Briefly, mice were anesthetized with Isoflurane (2-4% in oxygen, Abbott Laboratories, Green Oaks, IL), and mounted in a stereotaxic frame with a water-filled heating pad (38°C) beneath to maintain body temperature. An injection of a local anesthetic (0.05% marcaine, 0.1 ml s.c.) was administered before exposing the dorsal skull. A craniotomy was made to access the OT and pPCX, each contralateral from another (see **Fig S1A** for distribution). An 8-channel micro-wire electrode array (PFA-insulated tungsten wire, with four electrode wires encased together in a 254 µm diameter polyimide tube (Gadziola et al., 2015)) was implanted within the OT (1.25mm lateral, 1.25mm anterior bregma, 4.9mm ventral) and posterior PCX (3.9mm lateral, 1.5mm posterior bregma, 4.2 mm ventral) (**Fig 1A and S1A**), and cemented in place, along with a head-bar for later head fixation. The hemisphere receiving an array into each structure was not held constant across animals but instead varied (**Fig S1A**). Additional craniotomies were drilled over a single neocortex for placement of a stainless steel ground wire. During a three-day recovery period, animals received a daily injection of carprofen (5 mg/kg, s.c., Pfizer Animal Health, New York, NY) or meloxicam (5mg/kg; Putney, Inc., Portland, ME) and *ad libitum* access to food and water. Animals were allowed 5-7 days for post-op recovery before beginning water restriction.

#### Olfactory lick/no-lick odor discrimination and reversal learning behavioral task

Mice were mildly water-restricted for three days prior to behavioral training. Bodyweight was monitored daily and maintained at 85% of their original weight. Mice were trained in cohorts of three to four. All behavioral procedures were performed during the light hours, in a dim room. Head-fixed mice were trained in a lick/no-lick odor discrimination task (**Fig 1C**), involving 4 different odors (2 reinforced and 2 unreinforced), across multiple 1 hour recording sessions in which the mice obtained a fluid reward for licking a spout positioned in front of their snouts in trials with reinforced odors (Gadziola et al., 2015), adapted from (Verhagen et al., 2007).

Licking was measured by a pair of infrared photo-beams positioned to cross in front of the lick spout by ~2 mm. Mice were first trained to lick the water spout for reward, with a progressively increasing inter-trial interval (ITI) (3 ± 1 s, Phase 1). In Phase 2, odor presentation began with a 10 ± 2 s ITI, and mice were only rewarded for licking during a 2 s period after odor offset (FR1). Licking during the odor was discouraged by moving the lick spout further away from the snout. In Phase 3, trials were randomized between rewarded and unrewarded trials (17 ± 2 sec ITI); for unrewarded odor trials, mice were presented with a “blank” stimulus (mineral oil) and had to learn to withhold licking during these trials. Finally, in Phase 4, trials (17 ± 2 s ITI) were randomized between the rewarded and unrewarded odors. In go trials, mice would receive a reward for licking a spout within 2sec following the offset of the rewarded odor (hit); not licking would be considered a miss. In no-go trials, mice were presented with an unreinforced odor and did not receive a water reward regardless of whether they licked (false alarm) or correctly withheld licking for the total odor duration of the trial (correct reject). Next, mice continued on Phase 5 to learn a novel odor pair, consisting of a novel reinforced odor and a novel unreinforced odor. Phase 5 consisted of pseudo-random trials (17 ± 2 s ITI) among which all four odors (2 familiar and 2 novel) were separately presented, with the two reinforced odors both resulting in a low-value reward. Finally, in Phase 6, the four odors were presented (17 ± 2 s ITI), with one rewarded odor being assign to the high value saccharine and the other to the low value saccharine. Mice discriminated among the four odors, two of which predicted presentation of reward (either low or high value saccharine), and two unrewarded odors. These sessions include the original training odor set, as well as novel odors presented on a different experimental day (each mouse was shaped on 2 novel sets of four odors on different sessions). No major effects were observed between high versus low value rewarded odor evoked responses nor behavior and therefore herein any odors paired with reinforcers are classified as conditioned rewarded odors, whereas those not paired are termed conditioned unrewarded odors. Throughout all phases, behavioral performance was evaluated in blocks of 20 trials, and mice were required to achieve a performance criterion ≥80% correct for two consecutive blocks in order to advance to the next phase (**Fig S1B**). Neural activity was recorded throughout all training phases, but analysis of odor-evoked activity was restricted to post-training sessions.

#### Stimulus Delivery

Odors were presented through a custom air-dilution olfactometer with independent stimulus lines up to the point of entry into the odor port. In addition to a blank stimulus (mineral oil), odors included ethyl butyrate, 1,7-octadiene, isopentyl acetate, heptanal, 2-heptanone, (+)-limonene, ethyl propionate, (−)-limonene, methanol, methyl valerate, 2-butanone, 1,4-cineole, butanal, propyl acetate, allylbenzene, allyl bromide, isobutyl propionate, and 2-methylbutyraldehyde (Sigma Aldrich, St. Louis, MO; all >97% purity). These molecularly diverse odors were diluted in their liquid state to 1 Torr (133.32 Pa) in mineral oil and were then further diluted to 10% (vol/vol) by mixing 100 ml odor vaporized N2 with 900 ml medical grade N2 (Airgas, Radnor, PA). Thus, stimuli were delivered at a total flow rate of 1 L/min. Not all animals were tested with all odors. The rewarded and unrewarded odor pairs were pseudo-randomly assigned to each cohort prior to training (neurons were not initially screened for odor responsiveness). The experimenter was not blind to odor assignment, but all stimulus presentation was automated. Presentation of rewarded and unrewarded odors were pseudo-randomized within each block, delivered for 2 s duration with a 17 ± 2 s ITI through a Teflon odor-port (9 mm diameter opening) directed towards the animal’s snout at a distance of 1 cm. Odor was continuously flowing to the odor-port but was removed by a vacuum before exiting towards the animal. Recordings with a photoionization detector (miniPID, Aurora Scientific, Ontario, Canada) were used to confirm the temporal dynamics of the odor presentation in this design (**Fig 1D**). While the dynamics may vary slightly across odors, they confirm the precision and stability of the odor presentation methods used in this study.

#### Reward Delivery

Reward fluids were delivered through a custom 3D-printed polylactic acid lick spout, as reported previously (Gadziola and Wesson, 2016). Independent stimulus lines terminated onto 20G blunted needles that passed through one of seven 1-mm holes and extended to the tip of the spout. In the current task, two adjacent holes on the lick spout were used for reward delivery, three were connected to a vacuum line, and the last two unused holes were blocked. Animals were reinforced with a 4µL drop of 2-20mM saccharin (Sigma Aldrich, St. Louis, MO; dissolved in water). Reward volumes were calibrated for the individual reward valves.

#### In vivo Electrophysiology

The outputs of the electrode arrays were amplified, digitized at 24.4 kHz, filtered (bandpass 300-5000 Hz), and monitored (Tucker-Davis Technologies, Alachua, FL), along with licking (300 Hz sampling rate), and odor and reward presentation events. One electrode wire was selected to serve as a local reference for each headstage. Our electrode arrays were fixed in place and no attempt was made to record from unique populations of neurons on different sessions. To compensate for the possibility that the same neurons were recorded across multiple days, two different behavioral tasks were employed and statistical comparisons are only made within each task type. Sessions of the same task and odor pair were run for 1-3 consecutive days to achieve adequate behavioral performance and/or capture the dynamics of newly identified neurons. On average, 3.5 ± 1.9 single neurons were identified per mouse per session (range: 1-7 neurons), with an average of 1.4 ± 0.6 neurons recorded per viable electrode wire per mouse per session (range: 1-3 neurons). Numbers were comparable when separated by region (± 0.1 single units / region difference).

### Methods for fiber photometric-based imaging of D1- and D2-type dopamine receptor expressing OT neurons during lick/no-lick odor discrimination and reversal learning

#### Virus injection and optical fiber Implantation for fiber photometry

To achieve cell-type specific GCaMP6f (Chen et al., 2013) expression, D1- and D2-Cre mice were injected with AAV5-EF1a-DIO-GCaMP6f-WPRE (5.43^e13^ GC/ml, obtained from the Penn Vector Core in the Gene Therapy Program of the Univ of Pennsylvania). Stereotaxic surgery was performed as described above with exceptions as noted below. 1µl of virus was infused into the OT (1.25mm lateral, 1.25mm anterior bregma, 4.9mm ventral) at a rate of 0.1nl/sec and allowed to diffuse for 10mins before needle was slowly withdrawn over the subsequent 20mins. This injection paradigm was not intended to infect solely OT neurons, but to yield a bolus of infected cells which could be later targeted selectively for imaging by careful localization of the optical fiber. Following, the craniotomy was sealed with wax and the wound margin closed. Animals were allowed 2 weeks for the virus to transduce and then a 400µm core, 0.48NA optical fiber, threaded through metal ferrules were chronically implanted to terminate within the OT (same coordinates as above). Implants were cemented in place along with a headbar for later head fixation. As with the single-unit recordings, animals were allowed 5-7 days for post-op recovery before beginning water restriction. D1- and D2-Cre mice were tested in mixed cohorts.

#### Imaging

465nm (GCaMP excitation wavelength, driven at 210 Hz) and 405nm (UV excitation wavelength, driven at 330Hz; control channel) light emitting diodes were coupled to a 5 port fluorescence mini cube (FMC5, Doric Lenses) using 400µm core, 0.48NA, 2.5mm FCM optical fiber patchcords. Excitation and emission light were directed through a single optical fiber patchcord connected to the animal via a 2.5mm metal ferrule. Emission light was directed through the 5-port mini cube, and subsequently coupled to a pair of femtowatt photoreceivers (model 2151, Newport) for monitoring of the GFP and UV fluorescence. Photometry data were digitized and pulse demodulated at 1kHz, along with the timing of behavioral events (licking) and odor and reward delivery using a Tucker Davis Technologies digital processor (TDT). Parameters used for imaging and acquisition of photometry data were held constant across all mice.

#### Lick/no-lick odor discrimination and reversal learning during photometry

The methods for operant olfactory behavior employed for fiber photometry were mostly identical to those used for the single-unit recordings (see above) with two exceptions. First, during the fiber photometry imaging, mice were only asked to discriminate between two odors in any given session (one conditioned rewarded and one conditioned unrewarded). Second, instead of being shaped through phases 5 & 6 as described above for the mice contributing unit data, mice contributing photometry data were shaped in a reversal learning paradigm. For reversal learning, following reaching successful criterion performance in the original odor discrimination, the contingencies were reversed so that the original rewarded odor no longer signals reinforcement availability (now the ‘new unrewarded odor’) and the original unrewarded odor now signals reward availability (now the ‘new rewarded odor’).

### Methods for optogenetic stimulation of OT D1 expressing neurons and intra-cranial self-stimulation

#### Virus injection and optical fiber Implantation for opto-ICSS

The surgical methods to allow for expression of ChR2 in OT D1-expressing neurons are similar to those used for GCaMP6f as described above. Two surgical procedures were performed on each mouse. First, mice were intracranially injected with one of two floxed AAV vectors to allow for expression of AAV5.EF1a.DIO.hChR2(H134R).eYFP or mCherry (to yield optogenetic stimulation, 10^e12^ vg/ml) or AAV5.EF1a.DIO.eYFP or mCherry (as control, 10^e12^ vg/ml) using the OT injection coordinates as described above at 1µl volume. AAVs were obtained from the University of North Carolina Gene Therapy Vector Core. Second, mice were intracranially implanted with a unilateral optic fiber (300um, 0.39NA) into their OT (same coordinates as the injection) which was attached to a ferrule to allow light transmittance into the brain. AAV injections and ferrule implants occurred in the left hemisphere or right hemisphere to prevent biasing results towards effects that may be lateralized. Using a light meter (Thorlabs, Inc., Newton, New Jersey), all fiber optic implants were verified prior to implantation to deliver 6.5-7.5mW^3^ of light energy when using the same LED and LED driver as used during Opto-ICSS testing (see below).

#### Opto-ICSS apparatus and light stimulation

We used a dual nose-poke port apparatus wherein poke into each port is monitored by 880nm infrared photobeams and digitized for later analysis. The floor and three walls of the apparatus were made of black acrylonitrile butadiene styrene (ABS) plastic (30cm tall walls), consisting of a 15 x 15cm square floor and a guillotine style ABS door to allow gentle insertion and removal of the mouse from the testing apparatus. One wall of the apparatus was constructed of clear acrylic to allow visual monitoring of the mouse by an experimenter when needed. The ceiling was ‘open’ to allow for tethering of the animal to an optical rotary joint (Doric Lenses, Inc., Quebec, Canada) which was suspended above the apparatus. The nose-poke ports were 3D-printed out of polylactic acid plastic and were T-shaped with a 2cm opening which faced into the behavioral apparatus to allow nose-entry and monitoring of infrared beam breaks. Photobeam status (open or closed) was acquired continuously and relayed to a PC for data collection using an Arduino microcontroller board running custom code written in the Arduino open source language (Arduino, https://www.arduino.cc). Immediately next to each port was a 3mm white LED, also controlled by the Arduino, which could signal to the mouse active port status (on = active and optical stimulation available, off = inactive and no available optical stimulation).

Optogenetic stimulation (25Hz train, 15ms pulse width, 2 sec duration [for opto-ICSS the 2 sec stimulation occurred regardless of the nose-poke duration]) was accomplished by means of a 447.5nm LED (Luxeon Rebel ES, Luxeon Stars, Lethbridge, Alberta) driven by a Thorlabs T-Cube LED driver (Thorlabs) and controlled by the digital output of an Arduino microcontroller. This stimulation paradigm was selected following the results of physiological and behavioral pilot studies. An SMA adapter was adhered atop of the LED using optical epoxy to allow for butt coupling of an SMA terminated multimode fiber to the LED. The fiber used to connect the LED to the optic rotary joint as well as the fiber used to connect the rotary joint to the mouse were both 300µm core multimode fibers with a 0.39NA (same as the fiber used for the optical implants), encased in 2mm furcation tubing.

#### Opto-ICSS shaping and testing

All behavioral procedures were performed during the light hours, in a dim room. At the start of all sessions, the mice were first gently connected to the fiber optic tether. The mice were then placed into the apparatus and the door closed. The stages of shaping and testing in the opto-ICSS task are illustrated in **Figure 6** and occurred on separate daily 60min sessions. In phase 1, both ports were active (both active port LEDs were on) and nose-poke into either port triggered blue light stimulation. After performing ≥ 40 pokes into either or both ports in a given session, for at least two consecutive sessions, mice were then transitioned into phase 2 of the task. In phase 2, the port wherein the mouse displayed the greatest number of pokes on subsequent days, was set to be active (port LED on, blue light stimulation available) and the alternative port was inactive (port LED off, blue light stimulation unavailable). Only nose poke into the active port triggered light stimulation. After mice reached criterion performance in phase 2 (≥40 pokes, with ≥80% into the active port), they were switched to phase 3 wherein the active and inactive ports were reversed so that the previously inactive port is now the only port that would trigger light stimulation. Phase 3 is thus a test of reversal learning and ensures that the animal’s tendency to poke is indeed motivated by light-mediated ChR2 stimulation. Those mice that reached criterion performance on phase 3 (see above), were advanced onto phase 4. In all of the previous phases stimulation was available on an FR1 (fixed ratio of 1) schedule. In phase 4, both ports were rendered inactive (both port LEDs were off, blue light off) in a test of extinction over two sessions. Transitions between phases occurred on subsequent daily testing sessions. Mice that did not reach criterion performance on a given phase by the end of the 6^th^ session of that phase were eliminated from further testing.

### Histology

Following the end of experiments, mice were overdosed with Fatal-plus (0.01mL/g; Vortech Pharmaceutical, Dearborn, MI) and perfused with 10mL of cold saline followed by 15mL of cold 10% phosphate buffered formalin. Brains were stored in 10% formalin/30% sucrose (4ºC) prior to sectioning frozen at 40µm thickness on a Leica sliding microtome. Tissue was later mounted on slides using Fluoromount-G containing DAPI (for fluorescence analysis; 4’,6-diamidino-2-phenylindole; Invitrogen, Carlsbad, CA) or stained with 1% cresyl-violet (for tissue contributing single-unit data).

## QUANTIFICATION AND STATISTICAL ANALYSES

### Analysis of electrophysiology data

Single neurons were sorted offline in Spike2 (Cambridge Electronic Design, Cambridge, England), using a combination of template matching and cluster cutting based on principle component analysis. Single neurons were further defined as having <2% of the spikes occurring within a refractory period of 2 ms. Spike times associated with each trial were extracted and exported to MATLAB (Mathworks, Natick, MA) for further analysis. To examine modulations in firing rate within a single trial, spike density functions were calculated by convolving spike trains with a function resembling a postsynaptic potential (Thompson et al., 1996). Mean firing rates across trials were measured in 50 ms bins, along with the 95% confidence interval. Mean baseline firing rate for each neuron was averaged across a 2 s period prior to odor onset. Neurons were considered task-responsive if two consecutive bins within a 7 s period from odor onset were significantly different from the baseline rate, having non-overlapping confidence intervals. This liberal 7 s window size was chosen to capture any modulation in firing rate that may be related to odor presentation, licking behavior, and/or reward ingestion. To examine odor responses averaged across the population of neurons in both regions, we computed the change in firing rate at each time bin by subtracting the mean pre-stimulus baseline rate (calculated over the 2 s prior to stimulus onset) from the response rate. We also calculated Z-scored firing rates from each 50ms binned spike density function outcome (also with a −2sec prior to odor onset background window, with significance defined at +/- 2 SD).

Animals were typically presented with at least 40 trials of each rewarded and unrewarded odor set per session. To ensure that animals were engaged in the task, only blocks possessing high behavioral performance were analyzed. On some trials, mice may have been licking during odor presentation. Any trials in which the animal licked during the first 1.5s period after odor onset were removed. This ensured that any odor-evoked activity observed during the first several hundred ms was not due to licking-related activity. All 11 mice contributed usable single-unit data from at least one of the targeted locations. Of the 11 animals with bilateral implants, 10 were confirmed to reside within the OT and 8 within the pPCX. One of the 10 OT-confirmed implants did not yield well-isolated neurons and therefore did not contribute data to our analyses. From all mice contributing data, 7 yielded simultaneous OT and pPCX single-units.

#### Fano factor Analysis

Fano factor was used as an autocorrelation metric to characterize neural spiking variability as described in (Geisler and Albrecht, 1997; Miller, 2006; Shadlen and Newsome, 1998). Variability in the spiking of pPCX and OT neurons was evaluated by computing Fano factor values during Phase 4 of the lick/no-lick odor discrimination task. Fano factor values to examine background activity (−2000 to −500 ms before odor onset) and odor-evoked activity (0 to 1500 ms after odor onset) were evaluated separately. For each trial within these two windows, the mean spike count was divided by the variance, and all values were averaged within neurons.

#### Decoding Analysis

The Neural Decoding Toolbox (www.readout.info) (Meyers, 2013) was used to assess the decoding accuracy of pPCX and OT neuronal activity to predict the trial type during Phase 4 of the lick/no-lick odor discrimination task (hit, miss, false alarm, or correct reject). In MATLAB, odor onset-aligned spiking activity from pPCX and OT neurons were binned into sliding 150ms bins at a 50ms step size. To determine whether decoding accuracy improved as a function of pseudoensemble size, the pattern classifier was trained and tested on 10, 20, or 30 randomly selected neurons per region. First, a cross-validation procedure was used where data from 78 trials containing each of the 4 trial types were randomly selected to create a pseudoensemble of neurons. These pseudoensemble vectors were divided into 11 random splits (based on the maximum number of trial repetitions for each trial type that all neurons were subject to following elimination of early-lick trials). Each neuron’s data underwent z-score preprocessing to prevent favoring of neurons with higher firing rates by the pattern classifier. A pattern classifier was then trained on 10 of the 11 splits of data, and the remaining split was assigned as test data. The classification accuracy was determined based on zero-one-loss method, where the maximum correlation coefficient between the test and training data and expressed as a percentage of correct predictions. The whole procedure was repeated 500 times. The decoding accuracy for a null distribution was created from OT and pPCX data by repeating the above steps with trial labels that were randomly shuffled 20 times and tested 50 times per shuffle to yield 1000 points per brain region.

### Analysis of fiber photometry data

Off-line, the UV signal was subtracted from that of the GFP, filtered (2^nd^ order, 25Hz low-pass), smoothed (9 data-point moving average), and down sampled to 200Hz using a custom script written in Spike2 (Cambridge Electronic Design, Inc). Photometry recordings were normalized across trials off-line, using the mean signal across 3 s before odor delivery as a trial specific baseline for the ∆F/F values. Of the 7 D1-Cre animals with bilateral implants, 1 was confirmed to have poor/weak viral expression and one have a fiber tip residing outside of the OT. Of the 7 D2-Cre mice, 2 were confirmed to have fiber tips residing outside of the OT. Thus, in total, 5 mice per genotype, which had 1) criterion-level behavioral performance, 2) on target OT fiber tips, and 3) quality viral expression contributed data.

### Analysis of histological data

Microscopic examination of tissue was conducted on a Nikon Ti2e inverted fluorescent microscope equipped with 5MP color and 15MP monochrome cameras. Histological definition of brain regions was aided by use of a brain atlas (Paxinos and Franklin, 2000).

### Overall statistical methods

All performed tests were two-sided and met assumptions of normality (Kolmogorov-Smirnov test). Sample sizes are consistent with numbers reported in the field and no statistical method was used to predetermine these. Statistical analyses were performed in SPSS 22.0 (SPSS Inc., Chicago, IL) or MATLAB. All data are reported as mean ± SD unless otherwise noted. Specific details regarding single-unit and fiber photometry analyses can be found in above sections.

## DATA AND CODE AVAILABILITY

Reasonable requests for access to code generated for the collection, extraction, processing, and analysis of data contained herein can be made through email to the lead contact.

## References

Barnes, D.C., Hofacer, R.D., Zaman, A.R., Rennaker, R.L., and Wilson, D.A. (2008). Olfactory perceptual stability and discrimination. Nat Neurosci 11, 1378–1380.

Berridge, K.C. (2019). Affective valence in the brain: modules or modes? Nat. Rev. Neurosci. 20, 225–234.

Berridge, K.C., and Aldridge, J.W. (2008). Decision utility, the brain, and pursuit of hedonic goals. Soc. Cogn. 26, 621–646.

Calu, D.J., Roesch, M.R., Stalnaker, T.A., and Schoenbaum, G. (2007). Associative Encoding in Posterior Piriform Cortex during Odor Discrimination and Reversal Learning. Cereb Cortex 17, 1342–1349.

Carlezon, W.A., and Chartoff, E.H. (2007). Intracranial self-stimulation (ICSS) in rodents to study the neurobiology of motivation. Nat. Protoc. 2, 2987–2995.

Carlson, K.S., Gadziola, M.A., Dauster, E.S., and Wesson, D.W. (2018). Selective Attention Controls Olfactory Decisions and the Neural Encoding of Odors. Curr. Biol. 28, 2195–2205.e4.

Chapuis, J., and Wilson, D.A. (2011). Bidirectional plasticity of cortical pattern recognition and behavioral sensory acuity. Nat Neurosci 15, 155–161.

Chen, T.-W., Wardill, T.J., Sun, Y., Pulver, S.R., Renninger, S.L., Baohan, A., Schreiter, E.R., Kerr, R.A., Orger, M.B., Jayaraman, V., et al. (2013). Ultrasensitive fluorescent proteins for imaging neuronal activity. Nature 499, 295–300.

Chikazoe, J., Lee, D.H., Kriegeskorte, N., and Anderson, A.K. (2014). Population coding of affect across stimuli, modalities and individuals. Nat Neurosci 17, 1114–1122.

Dias, B.G., and Ressler, K.J. (2014). Parental olfactory experience influences behavior and neural structure in subsequent generations. Nat Neurosci 17, 89–96.

DiBenedictis, B.T., Olugbemi, A.O., Baum, M.J., and Cherry, J.A. (2014). 6-Hydroxydopamine lesions of the anteromedial ventral striatum impair opposite-sex urinary odor preference in female mice. Behav. Brain Res. 274, 243–247.

Doucette, W., Gire, D.H., Whitesell, J., Carmean, V., Lucero, M.T., and Restrepo, D. (2011). Associative Cortex Features in the First Olfactory Brain Relay Station. Neuron 69, 1176–1187.

Gadziola, M.A., and Wesson, D.W. (2016). The Neural Representation of Goal-Directed Actions and Outcomes in the Ventral Striatum’s Olfactory Tubercle. J. Neurosci. 36, 548–560.

Gadziola, M.A., Tylicki, K.A., Christian, D.L., and Wesson, D.W. (2015). The Olfactory Tubercle Encodes Odor Valence in Behaving Mice. J. Neurosci. 35, 4515–4527.

Geisler, W., and Albrecht, D. (1997). Visual cortex neurons in monkeys and cats: detection, discrimination, and identification. Vis Neurosci 14, 897–919.

Gire, D.H., Whitesell, J.D., Doucette, W., and Restrepo, D. (2013a). Information for decision-making and stimulus identification is multiplexed in sensory cortex. Nat. Neurosci. 16, 991–993.

Gire, D.H., Whitesell, J.D., Doucette, W., and Restrepo, D. (2013b). Information for decision-making and stimulus identification is multiplexed in sensory cortex. Nat Neurosci 16, 991–993.

Gong, S., Zheng, C., Doughty, M.L., Losos, K., Didkovsky, N., Schambra, U.B., Nowak, N.J., Joyner, A., Leblanc, G., Hatten, M.E., et al. (2003). A gene expression atlas of the central nervous system based on bacterial artificial chromosomes. Nature 425, 917–925.

Gore, F., Schwartz, E.C., Brangers, B.C., Aladi, S., Stujenske, J.M., Likhtik, E., Russo, M.J., Gordon, J.A., Salzman, C.D., and Axel, R. (2015). Neural Representations of Unconditioned Stimuli in Basolateral Amygdala Mediate Innate and Learned Responses. Cell 162, 134–145.

Gottfried, J.A. (2009). Olfaction and Its Pleasures: Human Neuroimaging Perspectives. In Pleasures of the Brain, M.L. Kringelbach, and K.C. Berridge, eds. (Oxford University Press), pp. 125–145.

Gottfried, J.A. (2010). Central mechanisms of odour object perception. Nat Rev Neurosci 11, 628–641.

Gunaydin, L.A., Grosenick, L., Finkelstein, J.C., Kauvar, I. V., Fenno, L.E., Adhikari, A., Lammel, S., Mirzabekov, J.J., Airan, R.D., Zalocusky, K.A., et al. (2014). Natural Neural Projection Dynamics Underlying Social Behavior. Cell 157, 1535–1551.

Heimer, L., Switzer, R.D., and Van Hoesen, G.W. (1982). Ventral striatum and ventral pallidum: Components of the motor system? Trends Neurosci. 5, 83–87.

Hickey, C., and Peelen, M.V. (2015). Neural Mechanisms of Incentive Salience in Naturalistic Human Vision. Neuron 85, 512–518.

Ikemoto, S. (2003). Involvement of the Olfactory Tubercle in Cocaine Reward: Intracranial Self-Administration Studies. J. Neurosci. 23, 9305–9311.

Ikemoto, S. (2007). Dopamine reward circuitry: Two projection systems from the ventral midbrain to the nucleus accumbens-olfactory tubercle complex. Brain Res Rev 56, 27–78.

Ilango, A., Kesner, A.J., Keller, K.L., Stuber, G.D., Bonci, A., and Ikemoto, S. (2014). Similar Roles of Substantia Nigra and Ventral Tegmental Dopamine Neurons in Reward and Aversion. J. Neurosci. 34, 817–822.

Isosaka, T., Matsuo, T., Yamaguchi, T., Funabiki, K., Nakanishi, S., Kobayakawa, R., and Kobayakawa, K. (2015). Htr2a-Expressing Cells in the Central Amygdala Control the Hierarchy between Innate and Learned Fear. Cell 163, 1153–1164.

Kass, M.D., Rosenthal, M.C., Pottackal, J., and McGann, J.P. (2013). Fear Learning Enhances Neural Responses to Threat-Predictive Sensory Stimuli. Science (80-.). 342, 1389–1392.

Kermen, F., Midroit, M., Kuczewski, N., Forest, J., Thevenet, M., Sacquet, J., Benetollo, C., Richard, M., Didier, A., and Mandairon, N. (2016). Topographical representation of odor hedonics in the olfactory bulb. Nat Neurosci 19, 876–878.

Knaden, M., and Hansson, B.S. (2014). Mapping odor valence in the brain of flies and mice. Curr. Opin. Neurobiol. 24, 34–38.

Kobayakawa, K., Kobayakawa, R., Matsumoto, H., Oka, Y., Imai, T., Ikawa, M., Okabe, M., Ikeda, T., Itohara, S., Kikusui, T., et al. (2007). Innate versus learned odour processing in the mouse olfactory bulb. Nature 450, 503–508.

Kudo, Y., Akita, K., Nakamura, T., Ogura, A., Makino, T., Tamagawa, A., Ozaki, K., and Miyakawa, A. (1992). A single optical fiber fluorometric device for measurement of intracellular Ca2+ concentration: Its application to hippocampal neurons in vitro and in vivo. Neuroscience 50, 619–625.

Kumar, S., von Kriegstein, K., Friston, K., and Griffiths, T.D. (2012). Features versus Feelings: Dissociable Representations of the Acoustic Features and Valence of Aversive Sounds. J. Neurosci. 32, 14184–14192.

Lebel, D., Grossman, Y., and Barkai, E. (2001). Olfactory learning modifies predisposition for long-term potentiation and long-term depression induction in the rat piriform (olfactory) cortex. Cereb. Cortex 11, 485–489.

Lewin, K. (1935). A Dynamic Theory of Personality (McGraw-Hill Book Company, Inc.).

Li, Q., and Liberles, S.D. (2015). Aversion and attraction through olfaction. Curr. Biol. 25, R120–R129.

Li, W., Howard, J.D., Parrish, T.B., and Gottfried, J.A. (2008). Aversive learning enhances perceptual and cortical discrimination of indiscriminable odor cues. Science (80-.). 319, 1842–1845.

Lobo, M.K., and Nestler, E.J. (2011). The Striatal Balancing Act in Drug Addiction: Distinct Roles of Direct and Indirect Pathway Medium Spiny Neurons. Front. Neuroanat. 5, 41.

Mandairon, N., and Linster, C. (2009). Odor perception and olfactory bulb plasticity in adult mammals. J. Neurophysiol. 101, 2204–2209.

Meyers, E.M. (2013). The neural decoding toolbox. Front. Neuroinform. 7, 8.

Miller, P. (2006). Analysis of Spike Statistics in Neuronal Systems with Continuous Attractors or Multiple, Discrete Attractor States. Neural Comput. 18, 1268–1317.

Millman, D.J., and Murthy, V.N. (2020). Rapid Learning of Odor–Value Association in the Olfactory Striatum. J. Neurosci. JN-RM-2604-19.

Morrison, S.E., and Salzman, C.D. (2009). The Convergence of Information about Rewarding and Aversive Stimuli in Single Neurons. J. Neurosci. 29, 11471–11483.

Murata, K., Kanno, M., Ieki, N., Mori, K., and Yamaguchi, M. (2015). Mapping of Learned Odor-Induced Motivated Behaviors in the Mouse Olfactory Tubercle. J. Neurosci. 35, 10581–10599.

Oettl, L.-L., Scheller, M., Wieland, S., Haag, F., Wolf, D., Loeb, C., Ravi, N., Durstewitz, D., Shusterman, R., Russo, E., et al. (2019). Phasic dopamine enhances the distinct decoding and perceived salience of stimuli. BioRxiv 771162.

De Olmos, J.S., and Heimer, L. (1999). The Concepts of the Ventral Striatopallidal System and Extended Amygdala. Ann. N. Y. Acad. Sci. 877, 1–32.

Paxinos, G., and Franklin, K. (2000). The Mouse Brain in Stereotaxic Coordinates (San Diego: Academic Press).

Pessoa, L., and Adolphs, R. (2010). Emotion processing and the amygdala: from a “low road” to “many roads” of evaluating biological significance. Nat Rev Neurosci 11, 773–783.

Roesch, M.R., Stalnaker, T.A., and Schoenbaum, G. (2007). Associative encoding in anterior piriform cortex versus orbitofrontal cortex during odor discrimination and reversal learning. Cereb Cortex 17, 643–652.

Ross, J.M., and Fletcher, M.L. (2018). Learning-Dependent and −Independent Enhancement of Mitral/Tufted Cell Glomerular Odor Responses Following Olfactory Fear Conditioning in Awake Mice. J. Neurosci.

Schoenbaum, G., and Eichenbaum, H. (1995). Information coding in the rodent prefrontal cortex. I. Single-neuron activity in orbitofrontal cortex compared with that in pyriform cortex. J Neurophysiol 74, 733–750.

Schoenbaum, G., Chiba, A.A., and Gallagher, M. (1998). Orbitofrontal cortex and basolateral amygdala encode expected outcomes during learning. Nat Neurosci 1, 155–159.

Schoenbaum, G., Chiba, A.A., and Gallagher, M. (2000). Changes in functional connectivity in orbitofrontal cortex and basolateral amygdala during learning and reversal training. J Neurosci 20, 5179–5189.

Schoenbaum, G., Setlow, B., Saddoris, M.P., and Gallagher, M. (2003). Encoding predicted outcome and acquired value in orbitofrontal cortex during cue sampling depends upon input from basolateral amygdala. Neuron 39, 855–867.

Shadlen, M.N., and Newsome, W.T. (1998). The Variable Discharge of Cortical Neurons: Implications for Connectivity, Computation, and Information Coding. J. Neurosci. 18, 3870–3896.

Soares-Cunha, C., Coimbra, B., Sousa, N., and Rodrigues, A.J. (2016). Reappraising striatal D1- and D2-neurons in reward and aversion. Neurosci. Biobehav. Rev. 68, 370–386.

Taverna, S., Ilijic, E., and Surmeier, D.J. (2008). Recurrent Collateral Connections of Striatal Medium Spiny Neurons Are Disrupted in Models of Parkinson’s Disease. J. Neurosci. 28, 5504–5512.

Thompson, K.G., Hanes, D.P., Bichot, N.P., and Schall, J.D. (1996). Perceptual and motor processing stages identified in the activity of macaque frontal eye field neurons during visual search. J. Neurophysiol. 76, 4040–4055.

Tolman, E. (1932). Purposive Behavior (New York: Appleton-Century).

Veldhuizen, M.G., Rudenga, K.J., and Small, D.M. (2009). The pleasure of taste, flavor, and food. In Pleasures of the Brain, M.L. Kringelbach, and K.C. Berridge, eds. (Oxford University Press), pp. 146–168.

Verhagen, J. V, Wesson, D.W., Netoff, T.I., White, J.A., and Wachowiak, M. (2007). Sniffing controls an adaptive filter of sensory input to the olfactory bulb. Nat Neurosci 10, 631–639.

Vicente, A.M., Galvão-Ferreira, P., Tecuapetla, F., and Costa, R.M. (2016). Direct and indirect dorsolateral striatum pathways reinforce different action strategies. Curr. Biol. 26, R267–R269.

Voorn, P., Jorritsma-Byham, B., Van Dijk, C., and Buijs, R.M. (1986). The dopaminergic innervation of the ventral striatum in the rat: A light- and electron-microscopical study with antibodies against dopamine. J. Comp. Neurol. 251, 84–99.

Wesson, D.W., and Wilson, D.A. (2011). Sniffing out the contributions of the olfactory tubercle to the sense of smell: hedonics, sensory integration, and more? Neurosci Biobehav Rev 35, 655–668.

White, K.A., Zhang, Y.-F., Zhang, Z., Bhattarai, J.P., Moberly, A.H., in ‘t Zandt, E., Peña-Bravo, J.I., Mi, H., Jia, X., Fuccillo, M. V., et al. (2019). Glutamatergic neurons in the piriform cortex influence the activity of D1 and D2-type receptor expressing olfactory tubercle neurons. J. Neurosci. 38, 9546–9559.

Wilson, D.A., and Sullivan, R.M. (2011). Cortical processing of odor objects. Neuron 72, 506–519.

Wilson, D.A., Sullivan, R.M., and Leon, M. (1987). Single-unit analysis of postnatal olfactory learning: modified olfactory bulb output response patterns to learned attractive odors. J Neurosci 7, 3154–3162.

Zhang, Z., Zhang, H., Wen, P., Zhu, X., Wang, L., Liu, Q., Wang, J., He, X., Wang, H., and Xu, F. (2017a). Whole-Brain Mapping of the Inputs and Outputs of the Medial Part of the Olfactory Tubercle. Front. Neural Circuits 11, 52.

Zhang, Z., Liu, Q., Wen, P., Zhang, J., Rao, X., Zhou, Z., Zhang, H., He, X., Li, J., Zhou, Z., et al. (2017b). Activation of the dopaminergic pathway from VTA to the medial olfactory tubercle generates odor-preference and reward. Elife 6, pii:e25423.

