## Supplemental Results for "A neural system that represents the association of odors with rewarded outcomes and promotes behavioral engagement"

<u>Contents:</u>

Supplemental Figure 1 and legend

Supplemental Figure 2 and legend

**Figure S1**

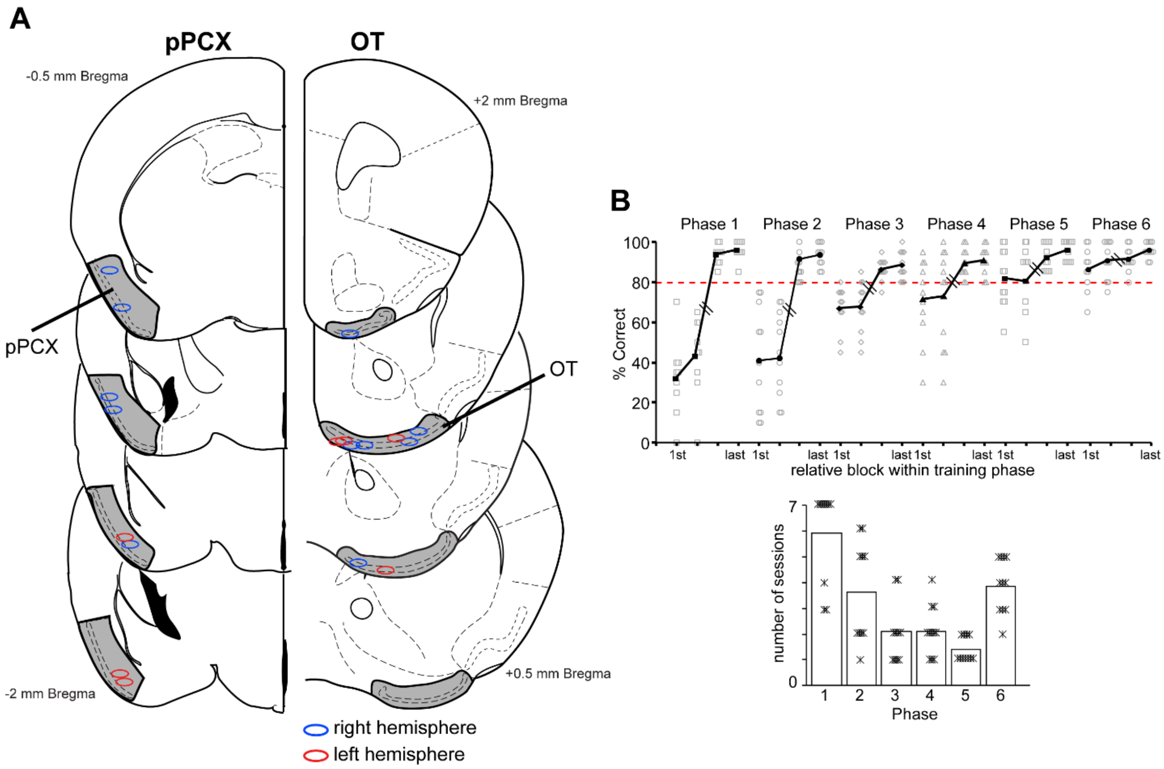


**Supplemental Figure 1. A)** Histological verification of pPCX and OT recording sites from mice contributing single-unit data. **B)** Summary of learning and performance in the lick/no-lick odor discrimination task for mice contributing single-unit data. Top panel displays the averages of behavioral performance in the first two and last two blocks of each of the six behavioral phases (see Methods). Each data point is from an individual mouse. Mice were transitioned onto the subsequent phase of testing after meeting the performance criterion (red dashed horizontal line, ≥ 80% correct responses) for at least two consecutive blocks. Chance performance = 50%. Lower panel indicates the average number of sessions to completion/criterion of each training phase. On average, following shaping, the mice used for analysis of single-unit data engaged in the odor discrimination task (across phases 4-6) with an average block performance of 92.6 ± 6.3% (mean ± SD).

### Figure S2

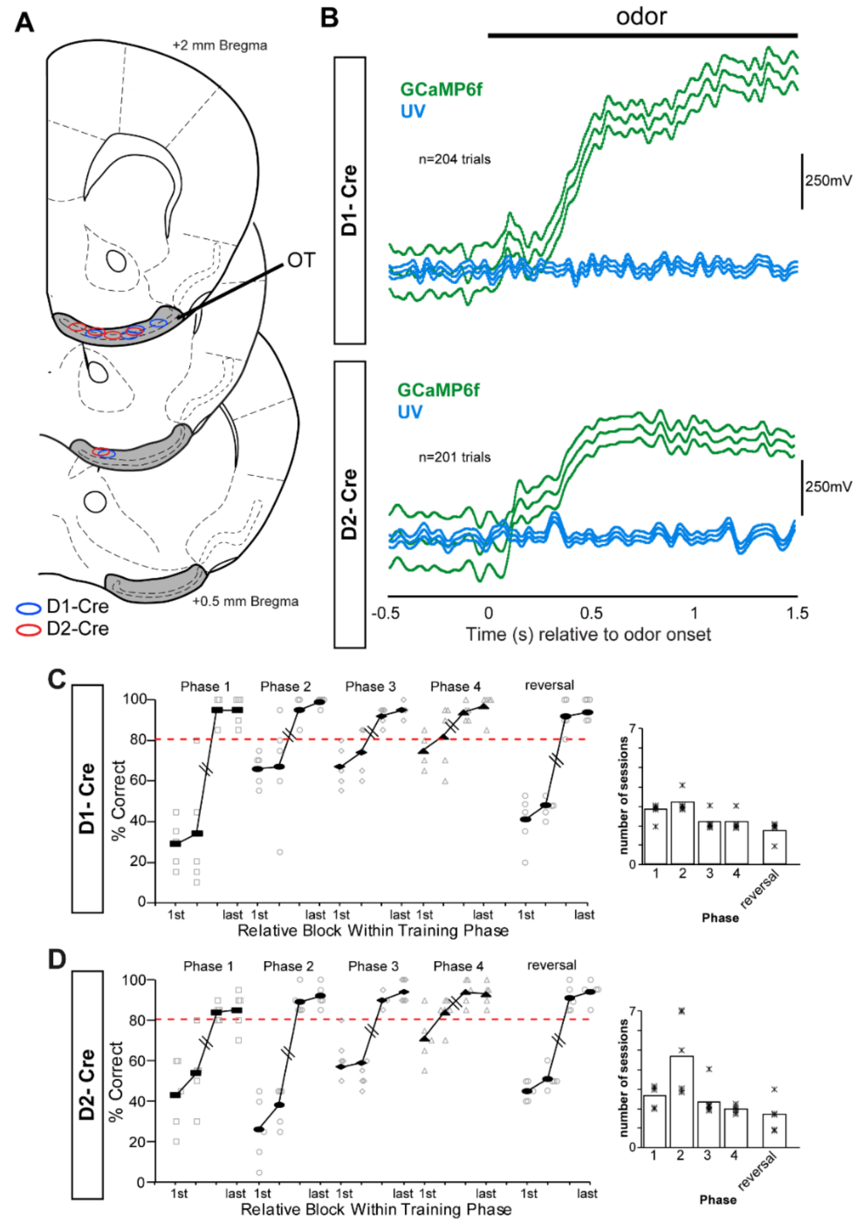


**Supplemental Figure 2. A)** Histological verification of optical fiber implants from mice contributing fiber photometry data. **B)** Example GCaMP6f and UV traces (2kHz sampling rate, 10Hz low-pass filter) averaged across trials of both conditioned rewarded and conditioned unrewarded odors throughout a single session from an individual D1-Cre (top) and D2-Cre mouse (bottom). These representative traces show the odor-evoked increase in GCaMP6f soon upon odor onset in contrast to the relative stability of the UV signal which would otherwise deviate from baseline in case of movement. All remaining photometry data included in this paper results from the GCaMP6f signal being subtracted from the UV signal for a single output, which is considered closely reflective of aggregate neural activity. Summary of learning and performance in the lick/no-lick odor discrimination and odor reversal learning task for D1-Cre **(C)** and D2-Cre mice **(D)** contributing fiber photometry data. Red dashed horizontal line = criterion behavioral performance level. Mice were transitioned onto the subsequent phase of testing after meeting the performance criterion (red dashed horizontal line) for at least two consecutive blocks. Chance performance = 50%. On average, following shaping, D1- and D2-Cre mice used for analysis of fiber photometry data engaged in the odor discrimination task (phase 4) with an average block performance of 93 ± 5.5% and 92.5 ± 5.5%, respectively (mean ± SD). Behavioral performance dropped towards chance for the first two blocks upon reversal learning which following shaping, reached criterion levels (≥ 80% correct responses for two consecutive blocks). Data from reversal sessions is an average across the numbers of odor pairs each mouse was conditioned among (range 2-3). Rightward panels indicate the average number of sessions to completion/criterion of each training phase.
